# Retinoid signaling promotes frontonasal identity while repressing maxillary identity during craniofacial development

**DOI:** 10.64898/2026.06.09.730988

**Authors:** Lin Xu, Yanran Wu, Maiko Omi-Sugihara, Takayuki Tsujimoto, Yujie Dang, Qi Wang, Xiuping Nie, Daisuke Motooka, Haruka Ohara, Toshihiro Inubushi, Masaya Yamaguchi, Lisa L. Sandell, Paul A Trainor, Takashi Yamashiro, Hiroshi Kurosaka

## Abstract

Vertebrate facial development depends on the correct specification of embryonic facial prominences, a process known to be governed by region-specific reciprocal signaling pathways between the craniofacial ectoderm, endoderm and mesenchyme. This process is further modulated by transcriptional and epigenetic mechanisms that regulate the expression of essential genes in mesenchymal progenitors derived from cranial neural crest cells. Retinoid signaling plays a critical role in facial development, and both gain-and loss-of-function results in a wide spectrum of facial defects, including orofacial clefts. In this study, we identified retinoid signaling as a critical regulator of cranial neural crest cell specification toward a frontonasal process identity in mice. Rdh10 oxidizes vitamin A (all-trans retinol) to retinal, which is a rate limiting step in the synthesis of retinoic acid. *Rdh10* loss-of-function resulted in ectopic formation of whisker pads - a derivative of the maxillary process - within the frontonasal process region. This transformation was evidenced by the mis-expression of maxillary specific transcription factors including *Meis2* and *Lhx6* in the frontonasal mesenchyme. Furthermore, these transcription factors exhibited increased chromatin accessibility at their consensus binding sites following the loss of retinoid signaling in frontonasal cranial neural crest cells. These results indicate that retinoid signaling acts as a critical regulator specifying frontonasal identity and fate of cranial neural crest cells as they migrate into the frontonasal process, while concomitantly repressing maxillary process fate. These results not only advance our understanding of frontonasal prominence specification and the evolutionary development of craniofacial structures but also offers valuable insights into the etiology and pathogenesis of craniofacial malformations such as orofacial clefts.

## Introduction

The process of facial development is highly conserved among vertebrates, and begins during early embryogenesis with the emergence of distinct facial prominences: the frontonasal process (FNP) anteriorly, a pair of maxillary processes (MxP) laterally, and a pair of mandibular processes (MnP) posteriorly (1, 2). These prominences are populated with mesenchymal cells largely derived from cranial neural crest cells (CNCCs), a progenitor cell population of varying potency that contributes to the majority of craniofacial tissues and structures (3–5). The correct acquisition of positional identity by CNCCs is crucial for proper craniofacial morphogenesis as CNCCs within distinct prominences give rise to specific structures. For example, CNCCs within the FNP contribute to the primary palate, nasal cartilage and upper incisors, whereas CNCCs within the MxP primarily generate osteogenic tissues and upper molars in the secondary palate (6, 7). In mice, CNCCs are induced at the neural plate border and migrate ventrolaterally to their craniofacial destinations by embryonic day (E)9.5 (3, 8). Throughout their migration and upon reaching their target regions, CNCCs continuously receive cues and guidance from various signaling pathways that shape their transcriptional networks and direct them toward appropriate craniofacial fates (9, 10). Disruption of these signaling pathways, can result in congenital anomalies such as mandibulofacial dysostosis (11) and auriculocondylar syndrome (12) in humans.

For example, *Endothelin 1* (*Edn1)* is expressed in the epithelium and mesodermal core of the MnP at E9.0 (13), and its deletion leads to transformation of the MnP to a MxP, and consequently mandible to maxilla (14). In contrast, ectopic expression of *Edn1* in CNCCs induces the transformation of the maxilla into a mandible (13). These studies also demonstrated that endothelin signaling is crucial for the induction of *Dlx5*, *Dlx6*, and *Hand2* expression in post-migratory CNCCs within the MnP to maintain their identity (14–16). Consistent with this model, *Dlx5/6* loss-of-function phenocopies the craniofacial anomalies characteristic of *Edn1-*null mice (14, 15). These results clearly demonstrate that specific signaling pathways in CNCCs are necessary and sufficient to determine their axial patterning and differentiated fates during facial development. Compared to this well-characterized molecular basis for maxillomandibular patterning and *Hox* gene regulation of more caudal pharyngeal arch patterning and development (17, 18), our understanding of what regulates the patterning and differentiation of the FNP remains relatively limited.

Retinoid signaling also plays a critical role in establishing the embryonic body plan, including the head and face (19). Retinoid signaling is mediated by all-trans retinoic acid (ATRA), the active metabolite of Vitamin A. The conversion of Vitamin A to ATRA is accomplished through two sequential oxidation reactions, with the first step, vitamin A (retinol) to retinal predominantly catalyzed during embryogenesis by retinol dehydrogenase 10 (RDH10) (20–22). The second oxidation of retinal to ATRA is performed by ALDH1A1, ALDH1A2 and ALDH1A3 (19, 23, 24). The active form of ATRA can either be degraded or function as a ligand to regulate gene expression at the transcription level by binding to heterodimers of nuclear retinoic acid receptors (RARs) and retinoid receptors (RXRs), which interact with retinoic acid response elements (RARE) in the promoter and enhancer regions of target genes (19, 25). In the embryonic craniofacial region, retinoid signaling is essential for the proper development of the maxillary complex, including the primary palate and secondary palate (20, 26–28). Perturbation of retinoid signaling leads to malformed midfacial structures and congenital craniofacial anomalies, such as orofacial cleft and choanal atresia through altered regulation of the *Alx* family of genes and Hh signaling. In this study, we found that active retinoid signaling is a critical determinant of CNCC specification within the FNP. By integrating single-cell transcriptomics and ATAC-seq analysis, we discovered a substantial molecular and epigenetic shift in CNCC fate from an FNP identity to a MxP identity upon the loss of retinoid signaling. Further analysis identified multiple transcription factors, including *Meis2*, *Lhx6* and *Alx1* that appear to be regulated by retinoid signaling and play central roles in driving this fate transition. These findings uncover novel mechanisms governing CNCC fate specification, particularly in the FNP, and provide critical insights into the molecular and epigenetic regulation of embryonic craniofacial development. Together with Endothelin-Dlx-Hand, and *Hox* gene mediated regulatory networks (11, 12, 15–18), RA-Alx signaling regulation of FNP patterning and development is a significant addition towards complete identification of the molecular repertoire that defines the regional identities of CNCC, as well as patterning the facial prominences and pharyngeal arches, and their generation of most of the craniofacial skeletons. This is important for a deeper understanding of the mechanisms that govern the development and evolution of the head and face, and for elucidating the basis of congenital craniofacial anomalies, which may inform therapeutic approaches.

## Results

### 1. Retinoid signaling is required for the development of the embryonic FNP

To investigate the role of retinoid signaling in regulating CNCC behaviors and embryonic craniofacial development, we analyzed the spatial and temporal distribution of retinoid signaling in the craniofacial region during the critical stages of CNCC induction, migration and colonization of the facial prominences and pharyngeal arches using *RARE-lacZ* reporter mice. Previous lineage tracing studies have demonstrated that CNCCs begin to delaminate from the dorsal neural plate boarder at the 4-somite stage (4ss), and actively migrate to initially colonize the facial prominences by the 8-10ss in mouse embryos (3, 8, 29, 30). At the 4ss and 9ss stages, we observed that retinoid signaling activity was present in the trunk of the embryo as expected, but was undetectable in the anterior neural plate border (induction site) (black arrowheads in Figure 1A, 1A’,1B and 1B’), or the midbrain and hindbrain regions (Figure 1A, 1A’, 1B, 1B’ and 1C), and the prospective FNP and first pharyngeal arch (asterisk in Figure 1B). Interestingly, a few hours of development later at the 14ss, which coincides with CNCC colonization of the FNP, we detected robust retinoid signaling activity specifically within the FNP in both facial ectodermal and mesenchymal cells, while it continued to be absent from the first pharyngeal arch (Figure 1C). Histological sections further revealed that retinoid signaling occurs in the mesenchyme cells of the FNP (Figure 1D). To assess the function of retinoid signaling, we genetically deleted *Rdh10* in a ubiquitous manner by administering tamoxifen to *R26-CreER^T2^;Rdh10^fl/fl^*embryos at E7.0, a condition under which we have previously confirmed near-complete elimination of *Rdh10* from the craniofacial region (27). Consequently, we observed a substantial reduction of retinoid signaling activity within the FNP of mutant embryos compared to controls (Figure 1E and 1E’).

**Figure 1:**
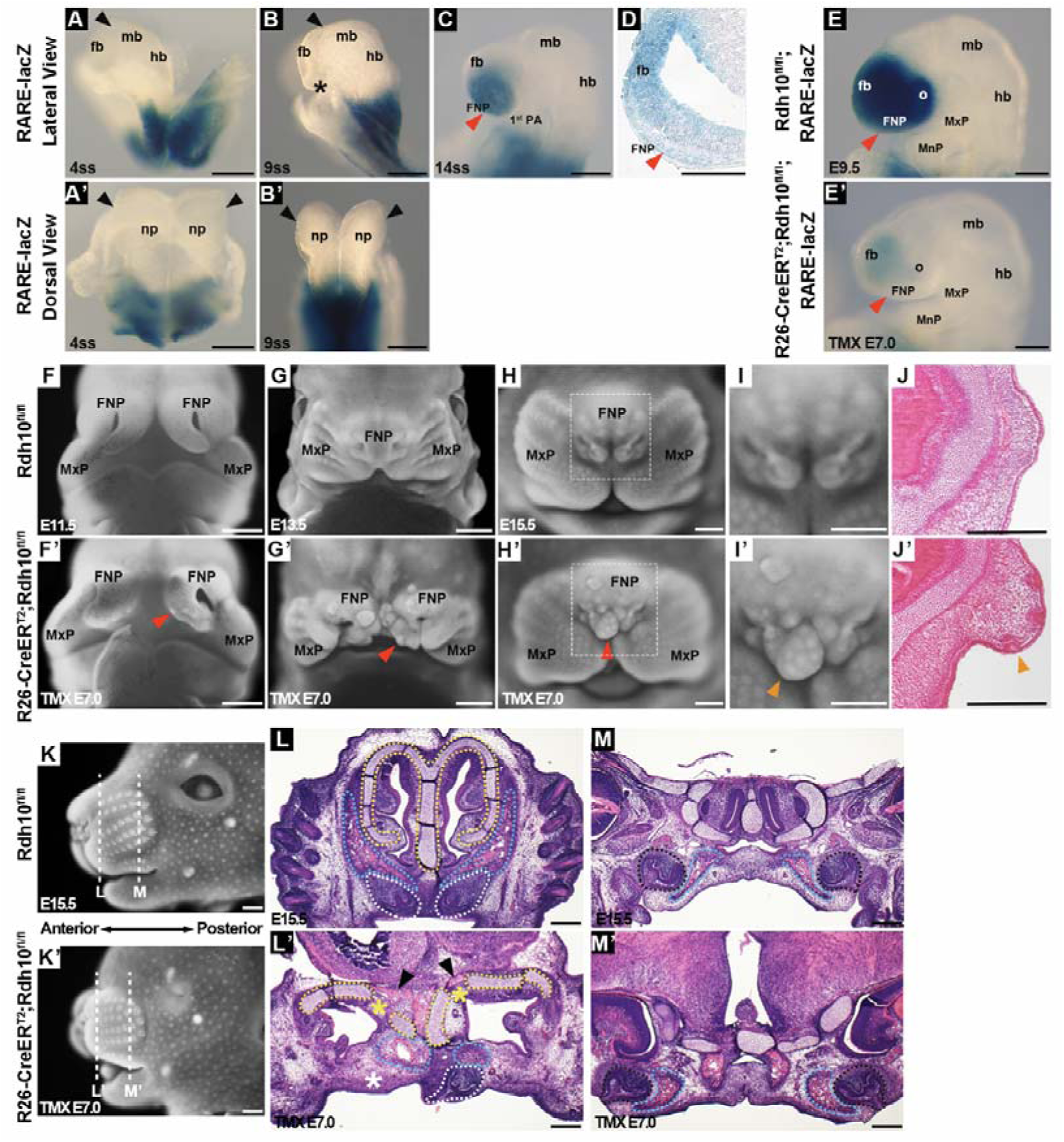
Retinoid signaling is required for the development of the FNP during embryogenesis. (A-E’) X-gal staining of *RARE-lacZ* embryos at 4ss (A, A’), 9ss (B, B’), 14ss (C). A-C show lateral views, while A’ and B’ show dorsal views. Black arrowheads in A-B’ indicate the CNCC induction site at the anterior neural plate border. The asterisk in B marks the location of the prospective FNP and first pharyngeal arch. Red arrowhead in C highlight the robust retinoid signaling in the FNP. (D) Representative sagittal sections of C. Red arrowhead in D indicate detectable β-galactosidase activity in the FNP, primarily localized to the mesenchyme. (E-E’) Lateral views of X-gal staining of E9.5 *R26-CreER^T2^;Rdh10^fl/fl^;RARE-lacZ* embryo (E’) which was treated with tamoxifen at E7.0 together with its littermate control (E). The red arrowhead in E’ highlights the diminished β-galactosidase activity in the FNP of mutant embryo. Scale bars, 200 μm. (F-J’) Frontal views of whole-mount DAPI staining in E11.5 (F-F’), E13.5 (G-G’) and E15.5 (H-H’) embryos. (I-I’) High-magnification images of the boxed regions in H-H’. (J-J’) Representative sagittal sections of H and H’. Red arrowheads in F’-H’ indicate ectopic protrusions. Orange arrowheads in I’-J’ highlight ectopic whisker pads forming within these protrusions. Scale bars, 500 μm. (K-M’) Lateral views of whole-mount DAPI staining in an E15.5 *R26-CreER^T2^;Rdh10^fl/fl^*embryo (K’) and its littermate control (K). White dashed lines in K and K’ indicate the planes used for sections shown in L-M’. (L-L’) Representative anterior sections, and (M-M’) representative posterior sections. Upper incisors are outlined with white dashed lines, upper molars with black dashed lines, nasal cartilage with yellow dashed lines, and maxillary bone with blue dashed lines. In L’, the white asterisk indicates the missing upper incisor, the yellow asterisk shows partially missing nasal cartilage, and the black arrowhead highlights ectopic bone formation. Scale bars, 500 μm. fb, forebrain; mb, midbrain; hb, hindbrain; np, neural plate; FNP, frontonasal process; 1^st^ PA, first pharyngeal arch; MxP, maxillary process; MnP, mandibular process; o, optic eminence.

We next analyzed the morphological outcomes of deficient retinoic acid signaling using whole-mount DAPI staining. At E11.5, we observed the formation of abnormal protrusions in the FNP of *R26-CreER^T2^;Rdh10^fl/fl^*embryos compared to controls (Figure 1F and F’). Notably, both the number and size of these protrusions progressively increased as development proceeded (Figure 1G and G’). By E15.5, ectopic whisker pads had developed within the abnormal protrusions of *R26-CreER^T2^;Rdh10^fl/fl^* embryos (Figure 1H-J and H’-J’). In addition, ectopic whisker pad formation was detectable only in mutant embryos administered tamoxifen at E7.0, whereas this phenotype became markedly less evident when tamoxifen was administered at E7.5 or later (Supplemental Figure 1). Histological sections further revealed significant defects in other craniofacial structures derived from the FNP such as absence of the upper incisor, partial loss of nasal septum and cartilage and ectopic bone formation in the nasal cartilage region (Figure 1K, K’, L and L’). In contrast, the upper molars and maxillary bone, which originate from the MxP, did not exhibit morphological abnormalities as severe as those observed in the FNP region (Figure 1M and M’). These findings demonstrate that temporally specific diminished retinoid signaling disrupts proper craniofacial patterning, resulting in the formation of ectopic protrusions and whisker pads, together with defects in the structures normally derived from the FNP.

### 2. Single-cell RNA sequencing analysis of FNP in *Rdh10* mutants

To uncover the molecular mechanisms underpinning the morphological effects of deficient retinoid signaling on FNP development, we performed single-cell RNA sequencing of frontonasal cells isolated from E11.5 *R26-CreER^T2^;Rdh10^fl/fl^* embryos and their littermate controls (Figure 2A). After applying the same quality filtering criteria to both datasets, we analyzed 9424 control cells and 7932 mutant cells. Uniform manifold approximation projection (UMAP) clustering of the two datasets combined revealed five major cell type populations, which were annotated based on the expression of known cell type-specific markers (Figure 2B and 2C). The identified clusters were consistent with the known In vivo cellular composition of the FNP and aligned well with previously published datasets (31, 32). To determine whether cell numbers across different populations were equally affected by the deficiency in retinoid signaling, we compared the proportions of cells in each cluster between control and mutant samples. A two-proportion z-test showed that the proportions of these five cell types were comparable between the two groups (Figure 2D), indicating that retinoid signaling deficiency does not significantly alter the overall cellular composition of the FNP at E11.5. We next assessed the impact of diminished retinoid signaling on transcriptional profiles within individual cell types by visualizing the UMAP plots for the control and mutant datasets separately. While most cell types displayed similar molecular profiles between the two groups, we observed a substantial difference in the size and distribution of the mesenchymal cell cluster in the mutant compared to control (Figure 2E and 2F). The single frontonasal mesenchymal cluster observed in controls was split into two distinct clusters in the mutants—one representing a reduced population of frontonasal mesenchyme, and the other, an ectopic mesenchymal cluster absent in controls (Figure 2F). These findings indicate that reduced retinoid signaling during frontonasal development specifically alters the transcriptional profile of mesenchymal cells, the majority of which are derived from CNCCs. This cellular reprogramming likely underlies the formation of ectopic structures observed in the FNP of *Rdh10* mutant embryos.

**Figure 2:**
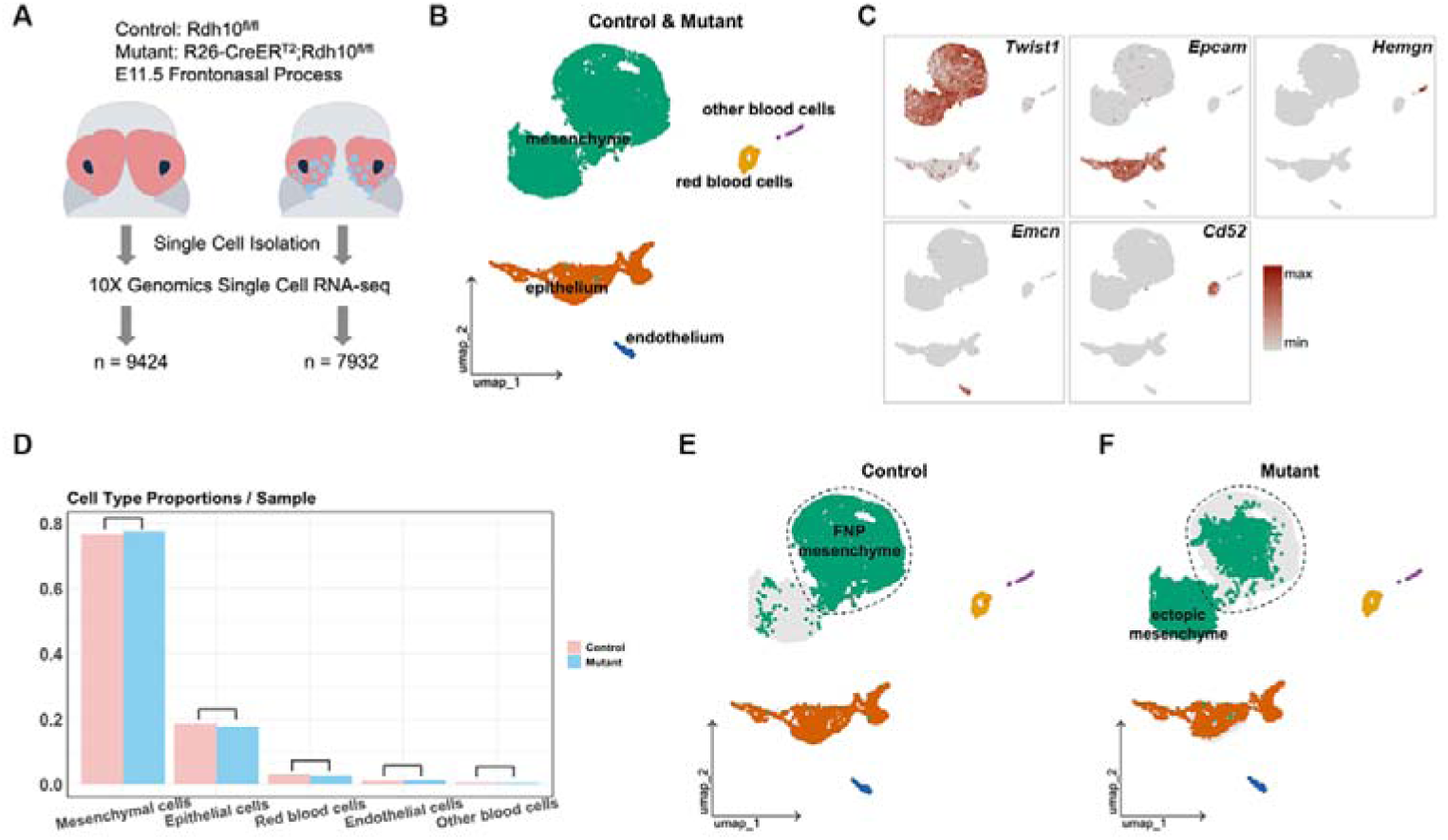
Comparable mesenchymal cell numbers but distinct transcriptomes between control and *Rdh10* mutants. (A) Workflow illustrating the process of single-cell RNA sequencing analysis. (B, C) UMAP visualization of distinct cell populations in control and *R26-CreER^T2^;Rdh10^fl/fl^* samples (B). Populations were annotated based on marker gene expression (C). (D) Bar chart comparing the percentage of distinct cell types between control and *R26-CreER^T2^;Rdh10^fl/fl^* samples based on single-cell RNA sequencing data. (E-F) UMAP comparisons of control (E), and mutant (F). In control (E), the FNP mesenchymal population is localized within a specific region circled by dashed line and labeled as “FNP mesenchyme” (E). In mutants (F), an additional mesenchymal population is present and labeled as “ectopic mesenchyme”

### 3. Loss of retinoid signaling converts caudal frontonasal CNCCs to a maxillary-like identity in *Rdh10* mutants

We next subsetted the mesenchymal cluster (7083 control cells and 6034 mutant cells) and re-clustered the data, which resolved into 4 distinct clusters (Figure 3A). Using cluster-specific markers, we identified three of these clusters representing cell populations from different anatomical locations: the caudal lateral nasal process (caudal LNP: *Pax7*), the rostral frontonasal process (rostral FNP: *Tfap2b*), and the caudal medial nasal process (caudal MNP: *Alx3*) (Figure 3A and 3B). These assignments were further validated by *in situ* hybridization (lower panel in Figure 3B). Notably, the fourth cluster, (denoted in yellow), did not express any of the frontonasal markers and was therefore annotated as “Ectopic non-FNP” (Figure 3A and 3B). By comparison, we observed a significant reduction of caudal MNP and caudal LNP cells in the mutant sample. In contrast, the majority of the ‘Ectopic non-FNP’ cluster was present in the mutant sample, while a small population was also included in the control samples, which is probably the result of minor contamination by anteriorly located MxP cells (Figure 3C and 3D). To define the expression profile of the diminished and ectopic mesenchymal populations, we performed unbiased differential transcriptome analysis between the control and mutant samples. The diminished cellular populations (caudal MNP and caudal LNP) exhibited high expression of *Rarb*, a known readout gene of retinoid signaling (33) (Figure 3E). Consistent with these results, we confirmed substantial downregulation of FNP marker genes, including *Pax7* and *Alx1* in the mutant FNP (Supplemental Figure 2). Feature plots of *Rarb* expression and in vivo *RARE-lacz* reporter results confirmed that ATRA target cells are highly enriched in the caudal MNP and caudal LNP regions of E11.5 control embryos with no noticeable overlap with *Rdh10* expression. In contrast this population was largely absent in the mutant (Figure 3F, Supplemental Figure 3). Conversely, the “Ectopic non-FNP” cluster predominantly expressed genes unique to the MxP, such as *Asb4*, *Lhx6* and *Lhx8* a tissue that was not included in this assay (highlighted in red in Figure 3G). Additionally, we did not detect activation of the Edn1–Dlx5/6–Hand2 signaling axis, which is characteristic of mandibular identity, in the ectopic non-FNP cluster (Supplemental Figure 4). These findings were validated through *in situ* hybridization, which demonstrated the ectopic expression of maxillary genes including *Asb4*, *Lhx6* and *Lhx8* in the developing FNP of mutant embryos (Figure 3H, Supplemental Figure 4). Further examination of stained embryos sections confirmed that the ectopic expression of *Lhx6* was specifically localized to the FNP mesenchyme (right panel in Figure 3H).

**Figure 3:**
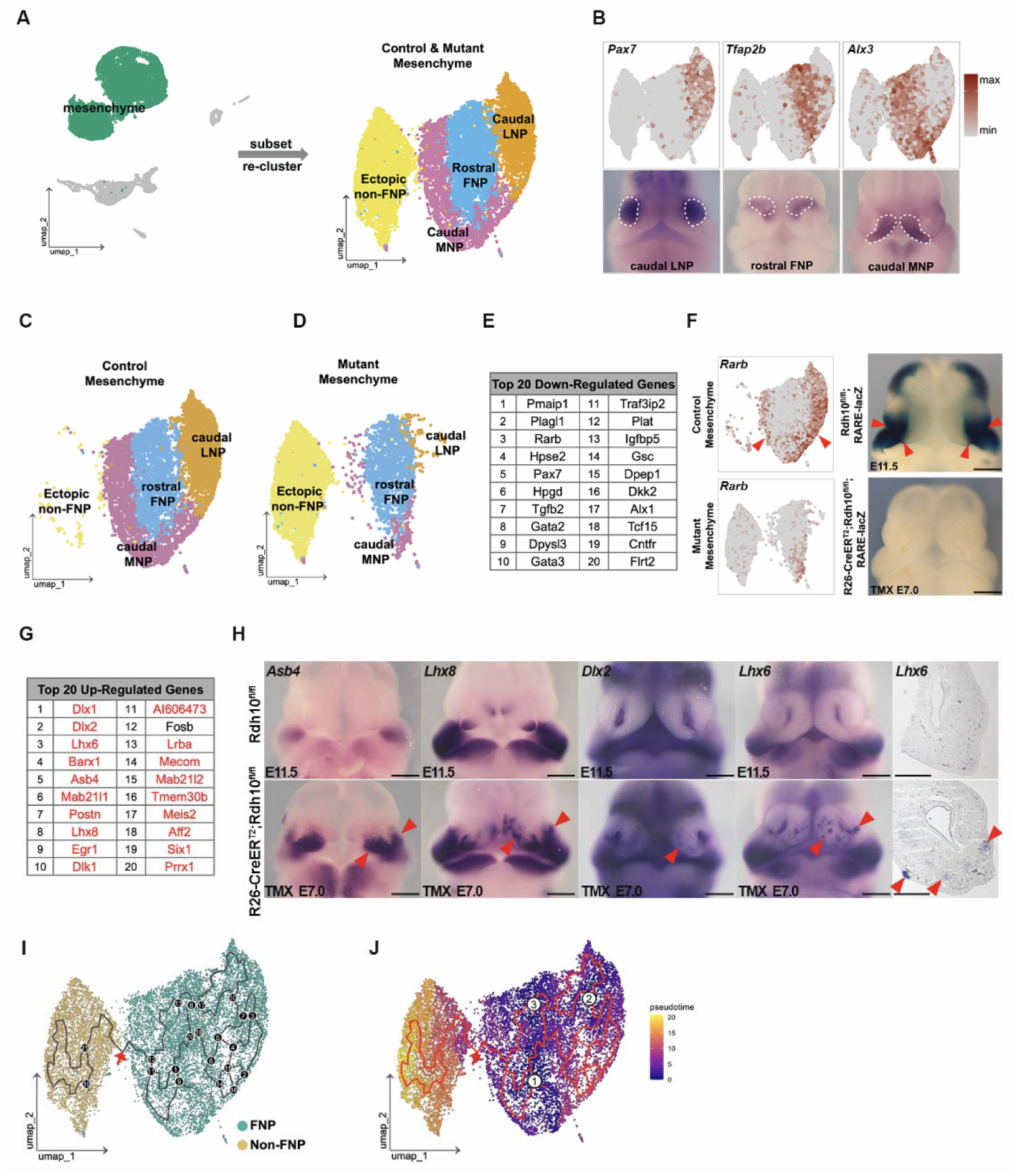
Diminished retinoid signaling converts caudal frontonasal CNCCs to a maxillary-like identity in *Rdh10* mutants. (A, B) UMAP visualization of re-clustered mesenchymal populations in *Rdh10^fl/fl^*and *R26-CreER^T2^;Rdh10^fl/fl^* samples (A). Populations were annotated based on known location-specific FNP marker expression (upper panel in B) and validated by whole-mount *in situ* hybridization using corresponding mRNA probes (lower panel in B). Caudal LNP, caudal lateral nasal process; Rostral FNP, rostral frontonasal process; Caudal MNP, caudal medial nasal process. (C-D) UMAP comparison of mesenchymal populations between controls (C) and mutants (D). (E, F) Table (E) lists the top 20 downregulated genes in mutant CNCCs compared to controls, with *Rarb* ranked third. Feature plot (F, upper left) shows *Rarb* expression in control, primarily localized in the caudal MNP and caudal LNP clusters (red arrowheads). In mutant (F, lower left), *Rarb*^+^ cells are highly reduced. Frontal view of X-gal staining in E11.5 *Rdh10^fl/fl^*;*RARE-lacZ* embryos (F, upper right) reveals ATRA-targeted cells predominantly in the caudal FNP (red arrowheads), which are undetectable in the mutant (F, lower right). Scale bars, 500 μm. (G, H) Table (G) lists the top 20 upregulated genes in the mutant CNCCs compared to control, with genes known to be expressed in the MxP highlighted in red. Whole-mount *in situ* hybridization (H) in E11.5 mutant embryos (lower panels) and controls (upper panels) shows expanded mRNA expression of select genes from Table G. Expanded expression regions are marked by red arrowheads. Representative frontal sections of the stained embryonic heads (H, right) demonstrate mesenchymal-specific ectopic expression (red arrowheads) in the mutant FNP. Scale bars, 500 μm. (I, J) Trajectory maps of control and mutant CNCCs illustrating cell fate decisions along branches. Branch nodes in (I) are indicated by black circles, representing points at which cells diverge toward alternative outcomes. Pseudotime is inferred in (J), with three root points in the FNP mesenchymal population labeled. The red arrow highlights the trajectory from the frontonasal CNCC population toward the ectopic non-FNP CNCC population.

To further validate these observations, we performed trajectory analysis to track potential cellular transitions within the dataset. The resulting trajectory map (Figure 3I and 3J) revealed a clear branch between the FNP and the ectopic cluster, suggesting that frontonasal CNCCs in mutant embryos undergo a transcriptional transformation toward a maxillary-like identity. These findings indicate that retinoid signaling is essential for maintaining the molecular identity of CNCCs within the FNP. In the absence of retinoid signaling, ATRA-targeted frontonasal cells undergo a shift toward a maxillary transcriptional profile, resulting in ectopic maxillary mesenchymal populations.

### 4. Single-cell RNA sequencing reveals largely overlapping transcriptome profiles between ectopic tissues and endogenous maxilla

Given the transcriptome heterogeneity observed at distinct positions within the MxP (34, 35), we sought to know how closely the transcriptome profile of these ectopic tissues in the mutant embryos align with that of the endogenous MxP. To address this question, we integrated an additional control dataset of maxillary complex that included both the MxP and the FNP (n=11116), providing a reference for the endogenous MxP transcriptome (Figure 4A). By pooling three datasets together and focusing on the mesenchymal populations, we detected two mesenchymal subclusters corresponding to the MxP and the FNP (Figure 4B).

**Figure 4:**
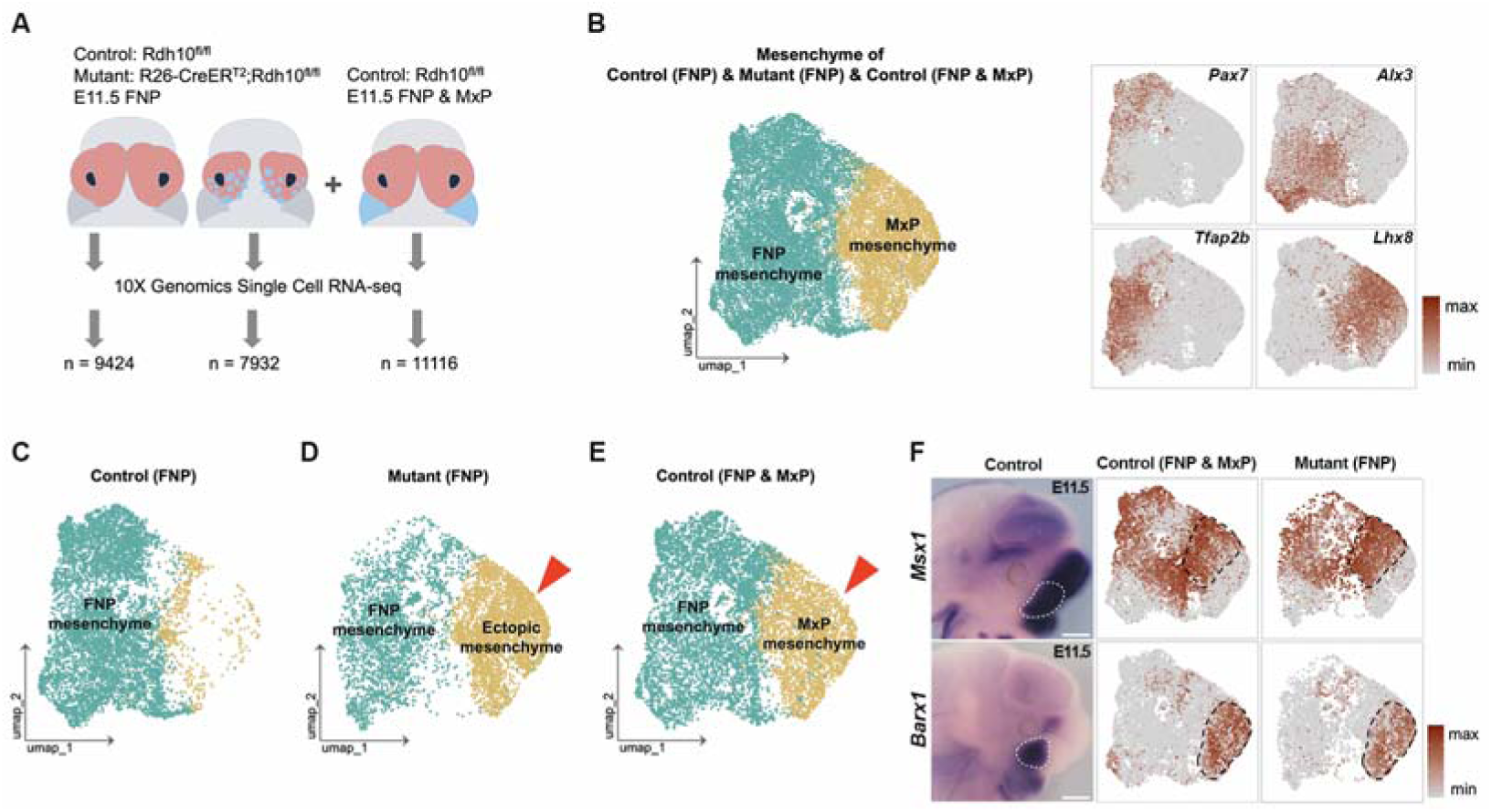
Single-cell RNA sequencing reveals identical transcriptome profiles between ectopic tissues in the mutant FNP region, and the control maxilla. (A) Workflow illustrating the single-cell RNA sequencing analysis process, which was integrated with another dataset using FNP and MxP tissues from E11.5 *Rdh10^fl/fl^* embryos. (B) UMAP visualization of the mesenchymal population including three samples, resolving into two subpopulations: the maxillary mesenchyme and the frontonasal mesenchyme. Feature plots (right panel) show the expression of marker genes used for annotation. (C-E) UMAP comparison of mesenchymal populations among three samples as indicated in the dataset accordingly. The red arrowheads indicate the ectopic mesenchyme in the mutant aligns closely with the natural maxillary mesenchyme. (F) Lateral views of *Msx1* and *Barx1* expression patterns in E11.5 control embryos (left panel). Expression domains within the MxP are outlined with white dashed lines. *Msx1* marks the anterior MxP, whereas *Barx1* marks the posterior MxP. Feature plots display *Msx1* and *Barx1* expression in control (middle panel) and mutant (right panel) samples. In the mutant, the ectopic cluster displays a gene expression profile comparable to the control maxillary cluster for both markers (black dashed lines), confirming its transcriptomic resemblance to the endogenous MxP. Scale bars, 500 μm.

Comparative analysis across all three datasets revealed a significant reduction in CNCCs in the FNP of *Rdh10* mutants which recapitulated our previous result (Figure 4D). Notably, the ectopic mesenchymal population in the mutant (Figure 4D) exhibited substantial transcriptional overlap with the endogenous MxP population from control embryos (Figure 4E). To further validate this overlap, we examined the expression of key MxP marker genes: *Msx1*, which is associated with the anterior MxP, and *Barx1*, which marks the posterior MxP (upper panel in Figure 4F). Feature plots revealed robust expression of both *Msx1* and *Barx1* within the ectopic mesenchymal cluster (lower panel in Figure 4F), suggesting that this population possesses a transcriptomic profile that overlaps with the endogenous MxP.

### 5. Activating retinoid signaling leads to the expression of FNP genes in the MxP

To test whether retinoid signaling is sufficient to reprogram CNCCs in the MxP into frontonasal identity, we treated embryos with ATRA dissolved in carrier or carrier alone as a control from the initial CNCC colonization stage (E8.5) (Figure 5A). To confirm the activation of retinoid signaling, we treated *RARE-lacZ* transgene reporter mice using this method and examined β-galactosidase activity. ATRA treatment resulted in robust activation of retinoid signaling across the entire embryo (Figure 5B’), including the first pharyngeal arch by E9.5 (red arrowhead in Figure 5C’). We then examined the positional identity of maxillary CNCCs by *in situ* hybridization using mRNA probes of *Alx1* and *Pax7* which usually mark the FNP. Compared to control, both *Alx1* and *Pax7* exhibited expanded expression domains that extended into the MxP in the ATRA-treated embryos (Figure 5D’, 5E’). These findings demonstrate that excessive retinoid signaling is sufficient to induce the ectopic expression of some FNP genes in CNCCs of the MxP.

**Figure 5:**
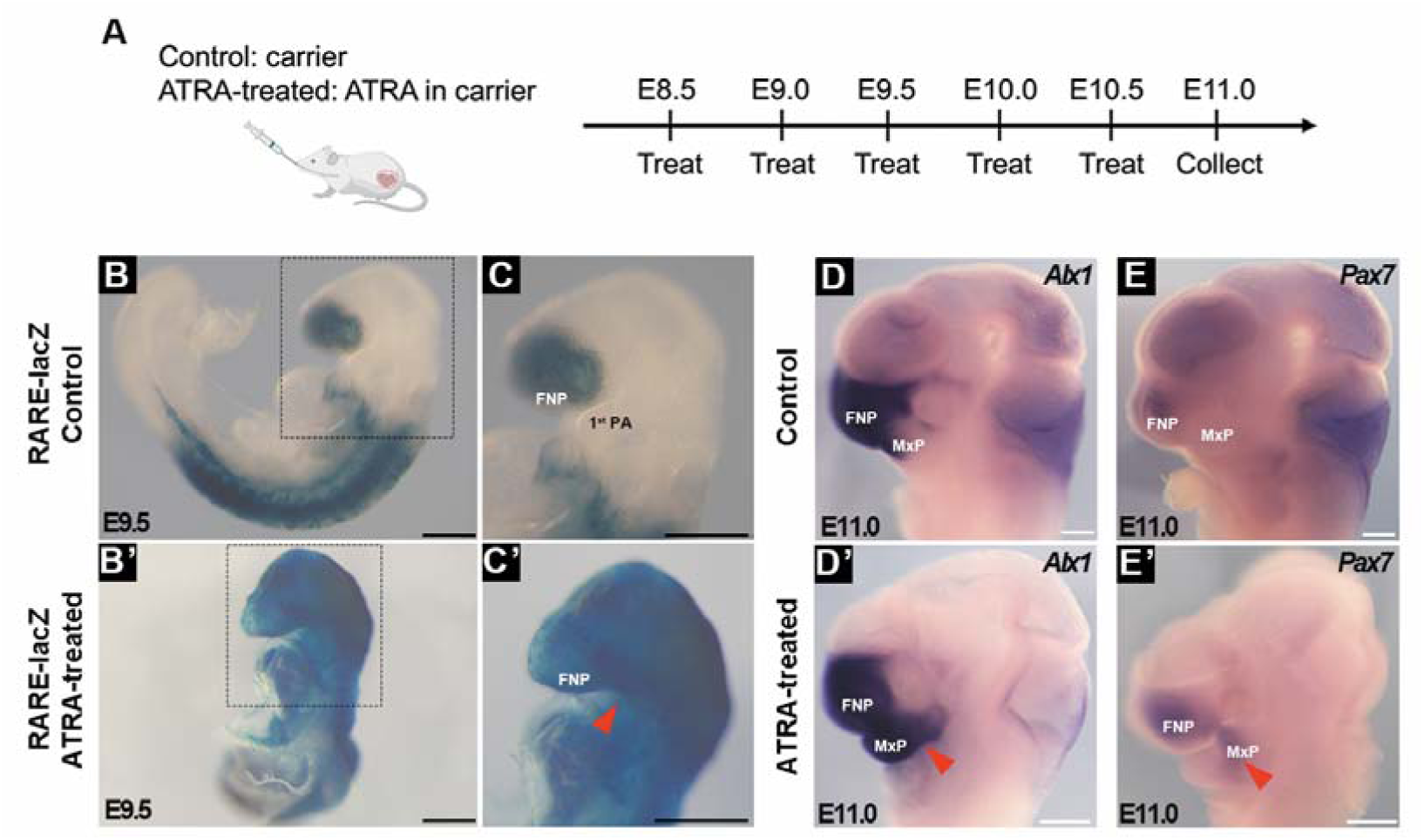
Mis-activation of retinoid signaling leads to the expression of frontonasal genes in the MxP. (A) Schematic drawing of the method to mis-activate retinoid signaling in the developing embryos created with BioRender.com. (B-C’) Lateral views of X-gal staining of E9.5 *RARE-lacZ* embryos, which were treated with control carrier (B) or ATRA with carrier (B’). C and C’ are the high magnification of the boxed areas in B and B’, respectively. The red arrowhead in C’ indicates robust retinoid signaling in the prospective first brachial arch. Scale bars, 500 μm. FNP, frontonasal process; 1^st^ PA, first pharyngeal arch. (D-E’) Lateral views of whole-mount *in situ* hybridization of E11.0 heads treated with control carrier (D,E) or ATRA with carrier (D’,E’) using indicated probes. The red arrowheads point to the ectopic expression of *Alx1* (D’) and *Pax7* (E’) in the MxP region. Scale bars, 500 μm. FNP, frontonasal process; MxP, maxillary process.

### 6. Single-cell multi-omics identifies key regulators in the transformed CNCCs

To further investigate the epigenetic mechanisms underlying the transcriptional alterations observed in the transformed CNCCs, we performed single-cell multi-ome sequencing (ATAC-seq and RNA-seq) of nuclei from frontonasal cells of E11.5 *R26-CreER^T2^;Rdh10^fl/fl^*embryos and controls (Figure 6A). After coarse clustering, we focused on the mesenchymal population which was resolved into two distinct subclusters: one corresponding to the ectopic MxP cells and the other representing the persistent rostral FNP CNCCs, as confirmed by the expression of *Lhx8* and *Tfap2b*, respectively (Figure 6B). Differential analysis of these two populations identified 26 transcription factors with significant enrichment in both RNA expression levels and chromatin accessibility at their motif regions in the ectopic MxP cells (log fold-change > 0.5, adjusted p-value < 0.05 for both RNA expression and motif enrichment) (Table S1). Dot plots highlighted the top 10 putative regulators, showing high enrichment of mRNA expression and motif accessibility specifically within the ectopic cell population (Figure 6C and 6D). To validate these findings, we performed Immunofluorescence to confirm the nuclear localization of the identified transcription factors. Notably, IHC results demonstrated that most of the identified transcription factors were co-localized in the nuclei of transformed caudal frontonasal CNCCs, including LHX6 and MEIS2 (Figure 6E). Interestingly, the regulatory region of Meis2 exhibited an anatomical position-dependent epigenetic status in control embryos, characterized by complementary enrichment of H3K27ac and H3K27me3, marks of active and repressive chromatin, respectively. Specifically, this region was marked by H3K27ac in the MxP, indicating an active chromatin state, while in the FNP, it was enriched for H3K27me3, reflecting transcriptional silencing (Supplemental Figure 5) (36). These findings strongly suggest that the transcription factors identified here, which are ectopically expressed in the transformed CNCCs, play central roles in mediating the switch in positional or regional identity of CNCCs from a frontonasal to a maxillary transcriptional program.

**Figure 6:**
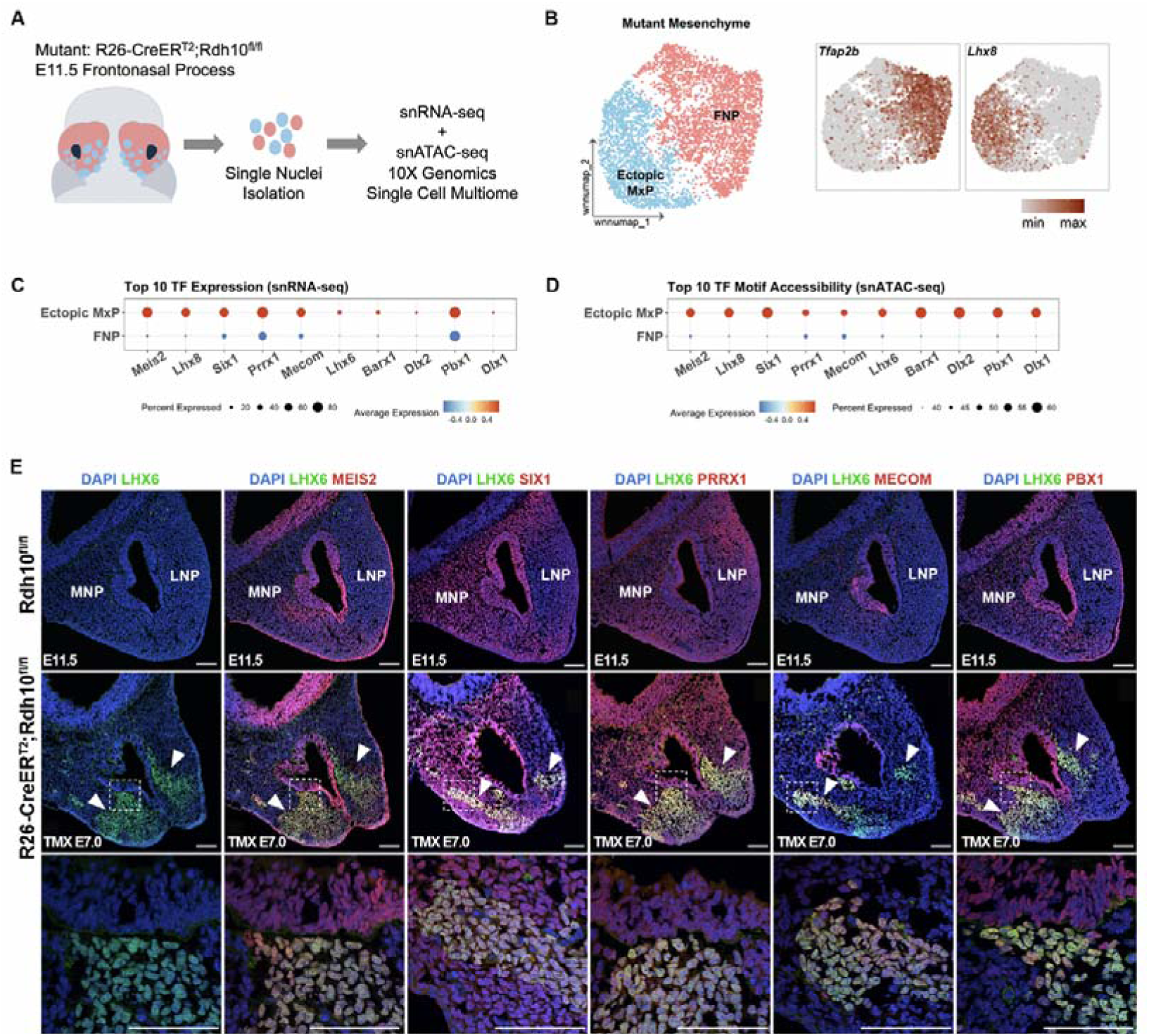
Single-cell multi-omics identifies key regulators in the transformed CNCCs. (A) Workflow illustrating the single-cell multiome analysis process, which performs RNA sequencing and ATAC sequencing of each single nuclei simultaneously. (B) UMAP visualization of mesenchymal populations resolved into two subclusters based on single-cell RNA sequencing and single-cell ATAC sequencing. The marker genes expressied in each cluster are shown on the panel on the right. (C-D) The mRNA expression level and motif accessibility in each cluster of the top ten transcription factors. (E) Frontal sections from E11.5 control (upper panel) and mutant (middle panel) embryos showing the distribution of the indicated protein (stained in red or green color) through immunofluorescence. DAPI counterstaining was used to visualize the cellular nuclei, which are shown in blue. The lower panel presents a high-magnification view of the boxed region. White arrowheads indicate the ectopic localization of maxillary marker proteins. Scale bars, 100 μm. MNP, medial nasal process; LNP, lateral nasal process.

## Discussion

During embryonic development, each facial prominence gives rise to specific craniofacial structures. For example, lineage tracing in mouse embryos has shown that the FNP gives rise to the medial and lateral nasal processes, with the medial nasal process contributing to the primary palate and nasal cartilage (6), while the lateral nasal process contributes to nasal cartilage (37). In contrast, the MxP forms structures such as the secondary palate and the whisker pad, a feature unique to the MxP, without contribution from other processes (6, 7). Analysis of retinoid activity during early craniofacial development using the *RARE-lacZ* reporter indicates that anterior CNCCs do not robustly receive retinoid signaling during their formation or migration, but rather only after they have reached and populated the FNP.

Functional analyses of retinoid signaling during embryonic craniofacial development have demonstrated its essential role in FNP formation. Genetic ablation of key enzymes required for all-trans retinoic acid synthesis, such as *Aldh1a2* and *Aldh1a3* results in severely disrupted FNP development in mouse embryos (38, 39). Similarly, studies in chick embryos have shown that early chemical elimination of retinoid signaling leads to profound frontonasal defects, whereas later perturbation results in milder phenotypes, highlighting the strong temporal dependence of RA signaling during facial morphogenesis (40). In addition, our group has previously demonstrated that reduced retinoid signaling perturbs midfacial structures, at least in part through aberrant elevation of Shh signaling (20, 26). Importantly, we have also previously reported that the overall phenotype of *Rdh10* mutant embryos, including craniofacial defects, can be largely rescued by administration of retinoic acid or retinal between E7.5 and E11.0 (20, 27). In addition, deletion of *Rdh10* at later stages than E7.5 produces considerably milder phenotypes (27). Together, these findings illustrate the tightly regulated spatiotemporal requirement for retinoid signaling during craniofacial development, with a critical window of effect between approximately E7.5 and E11.0 that overlaps with the period of craniofacial process specification.

In the present study, analysis of E15.5 *Rdh10* mutant embryos revealed ectopic whisker pad formation within the FNP, accompanied by the absence of structures such as the upper incisors and nasal cartilage. Ectopic structure formation during embryonic development occurs when cells undergo a switch in their developmental program (41, 42). Single-cell transcriptome and *in situ* hybridization analyses of the FNP of *R26-CreER^T2^;Rdh10^fl/fl^*embryos revealed a distinct ectopic cell population with a gene expression profile resembling that of the developing MxP. Notably, mesenchymal CNCCs underwent pronounced reprogramming in their gene expression profile, reflecting an FNP to MxP identity transformation. Importantly, we did not detect activation of the Edn1–Dlx5/6–Hand2 signaling axis in the ectopic non-FNP cluster which indicates the fate change is toward an MxP identity rather than a global pharyngeal segment transformation.

It is well established that retinoid signaling regulates a wide range of gene expression programs through transcriptional control mediated by retinoic acid receptors, which likely underlies the transcriptional changes observed in the present study (43, 44). We also identified that the cell population within the FNP that undergoes transformation corresponds to a caudal FNP CNCC population expressing genes such as *Rarb,* which is a nuclear receptor that binds RA to regulate gene expression, as well as *Pax7* and *Alx3*, whereas rostral FNP cells maintain their molecular identity even after elimination of retinoid signaling. These results indicate that not all, but rather a specific caudal subset of FNP CNCCs which are retinoid signaling responsive, transform to a maxillary prominence identity following the loss of retinoid signaling during the period of facial process fate determination in mice. Interestingly, *Alx3* loss-of-function in mice also resulted in partial expansion of MxP genes in the FNP (45). Additionally, mutation of the *alx* family of genes in zebrafish also resulted in transformation of FNP cells to a first branchial arch identity and subsequently a maxillary fate (Mitchell et al. Co-submitted paper). Collectively these results reveal there is an evolutionally conserved RA-ALX mediated molecular program that determines the regional identify and fate of FNP CNCCs during embryonic craniofacial development.

CNCCs comprise a progenitor cell population with varying degrees of potency that originate in the neuroepithelium, undergo an epithelial-to-mesenchymal transition, and migrate into the facial prominences where they develop into various craniofacial structures (4, 46). Neural crest cell patterning is regulated by a combination of signals they receive in the neuropeithelium during their development together with signals they receive from the ectoderm, mesoderm and endoderm environments during their migration and differentiation (17, 18, 47). Heterotopically transplanted CNCCs in mouse (48, 49), avians (50–52) and zebrafish (53) embryos, downregulate the molecular identity characteristic of their axial origin and adopt an identity appropriate for their new location (47). Neural crest cells are therefore inherently plastic and adaptable, and considered the conduit for much of craniofacial variation and evolution (54, 55). In our results, reduced retinoid signaling in the developing FNP resulted in alterations in the transcriptional profile of CNCCs from an FNP character to that of the MxP. Given that there is no significant retinoid signaling activity in the midbrain or adjacent mesenchyme — the regions where CNCCs that contribute to the FNP originate and initially migrate through — the molecular switch driving this plasticity is the reduced retinoid signaling encountered as the CNCCs reach the FNP.

Cellular transformation in vivo has been well-documented in model organisms. For example, in Drosophila, disruptions in the expression of the transcription factor Antp lead to head and thoracic transformations, demonstrating the powerful impact of transcriptional regulation on cellular identity (56). Similarly, cellular reprogramming often involves altered transcriptional regulation, as seen in the creation of induced pluripotent stem cells, where cells are reprogrammed to an undifferentiated state (57).

Underlying these processes, chromatin status plays a crucial role in allowing transcriptional regulation to function effectively. This is especially relevant in the craniofacial region, where epigenetic modifications of post-migratory CNCCs are essential for determining cell fate across various craniofacial prominences, guiding cells to adopt specific identities for different facial structures (36). Interestingly, our multi-ome analysis revealed several maxillary specific transcription factors were upregulated at the mRNA transcription level and this coincided with a high peak at consensus binding sequences for Meis2, Lhx8, Six1 and Prrx1. Notably, loss-of-function mutations in these factors are associated with craniofacial defects (58–61). A synergistic role for these ectopically expressed transcription factors in the FNP is seemingly responsible for transforming the cells’ transcriptional profile into one characteristic of the MxP. Additionally, *Meis2* has an ATRA-responsive sequence in its regulatory region consistent with being a direct target of RA signialing (62). Furthermore, CHIP-seq data for the active histone mark H3K27ac and the repressive mark H3K27me3 in the promoter region of *Meis2* in the FNP and MxP showed complementary patterns: H3K27ac exhibited a higher peak in the MxP, while H3K27me3 showed a higher peak in the FNP (Figure S1).

Importantly, retinoid signaling has been shown to regulate downstream transcriptional activity by recruiting histone modifiers around the RARE region, which plays a crucial role in controlling gene expression (62, 63). When considered alongside our findings, this suggests that retinoid signaling in the developing FNP modifies the regulatory regions of maxillary genes in a repressive manner. Consequently, reduced retinoid signaling lifts the repression on genes such as *Meis2*, *Lhx6*, *Lhx8*, *Six1* and *Prrx1* leading to a shift in the transcriptional profile of CNCC in the FNP, which ultimately changes their cellular identity and fate. In contrast, rostral FNP cells, which are largely devoid of retinoid signaling, retain their molecular identity even in *Rdh10* mutants.

A similar effect has been observed with respect to Fgf8 signaling in axial patterning during embryogenesis in response to diminished RA signaling (64). Directly related to our findings in this study, zebrafish frontonasal CNCC are also transformed to a dorsal first branchial arch identity when multiple *alx* genes are mutagenized. The further supports our model that the transformation in patterning and regional identity as a consequence of the loss of RA signaling is due to changes in *Alx* transcription factor expression (Mitchell et al. Co-submitted paper). It will be important in the future to validate the direct regulation by retinoid signaling through chromatin immunoprecipitation experiments using antibodies against retinoic acid receptors.

Interestingly, a previous study showed that enhancing retinoid signaling while inhibiting BMP signaling in the MxP of chick embryos induces formation of an ectopic egg tooth—a structure usually derived from the FNP (65). Our findings further demonstrate that excessive retinoid signaling expands the expression of frontonasal genes including *Pax7* into the MxP, indicating that retinoid signaling positively regulates frontonasal gene expression. At the same time, this retinoid signaling appears to repress maxillary gene activity, helping to maintain the distinct identities of the FNP and MxP. Together, these results highlight the crucial role of retinoid signaling in craniofacial patterning by promoting frontonasal-specific genes while inhibiting maxillary gene expression, thus guiding CNCCs toward their respective fates.

The determination of the identity of the maxillary and MnP within the first pharyngeal arch has been relatively well studied and is known to involve the endothelin signaling pathway, with *Dlx5* and *Dlx6* acting as downstream targets of this signaling. In contrast, the mechanisms responsible for determining the identity of the FNP and its interaction with the MxP remain an area of active research. Despite some progress, the precise regulatory networks guiding these processes are still being explored, highlighting the complexity of craniofacial development. Previous research in mice has demonstrated ectopic expression of maxillary genes in the FNP in association with mutations in *Alx* genes, which play a central role in frontonasal development (45). However, this is the first study to show ectopic formation of organs derived from the MxP, such as the whisker pad, in the FNP, together with clear evidence of a transcriptional profile switch at the single cell level. Importantly, we also observed the converse phenomenon in the expression of several genes between the FNP and MxP when exposed to excessive retinoid signaling. This study, in conjunction with previous findings, reveals the mechanisms underpinning the critical role of retinoid signaling in guiding the fate decisions of post-migratory cranial neural crest cells into FNP mesenchyme. Moreover, our results suggest that retinoid signaling acts as a key regulator, both repressing maxillary genes and activating frontonasal genes during early facial development.

It has been demonstrated that specific craniofacial prominences arose developmentally at distinct points during the evolutionary process. For example, the mandible, a derivative of the first pharyngeal arch in gnathostomes, is thought to have arisen through a heterotopic shift in epithelial-mesenchymal interactions mediated by facial epithelium and cranial neural crest cells (66). Intriguingly, temporally specific pulses of retinoid signaling occurring after cranial neural crest cell migration have also been shown to influence maxillary and mandibular fate determination in diverse vertebrates, including mice and frogs. In particular, suppression of retinoid signaling in the first pharyngeal arch appears necessary for proper developmental programming of the MxP and MnP, potentially recapitulating the process of heterotopic shifts in gene expression patterns that lead to the acquisition of distinct identities for different facial processes (67, 68). Unlike the first pharyngeal arch, our results showed that retinoid signaling received after the completion of rostral CNCC migration around E9.0 is critical for the proper development of the FNP during embryogenesis. These results suggest that the differentiation of CNCCs in specific facial prominences involves distinct sets of signaling pathways and transcription factors. *Hox* genes govern the patterning and regional identities of the caudal pharyngeal arches (17, 18), whereas Endothelin-Dlx-Hand signaling regulates maxillomandibular patterning and identity (11, 12, 15, 16). Our data demonstrates that retinoic acid signaling potentially acts as a central determinant of FNP patterning and identity (Figure 7). This is an important and significant step towards full resolution of the molecular repertoire defining the regional identities of CNCC, as well as patterning the facial prominences, the pharyngeal arches and their cranioskeletal derivatives, during vertebrate evolution and development.

**Figure 7:**
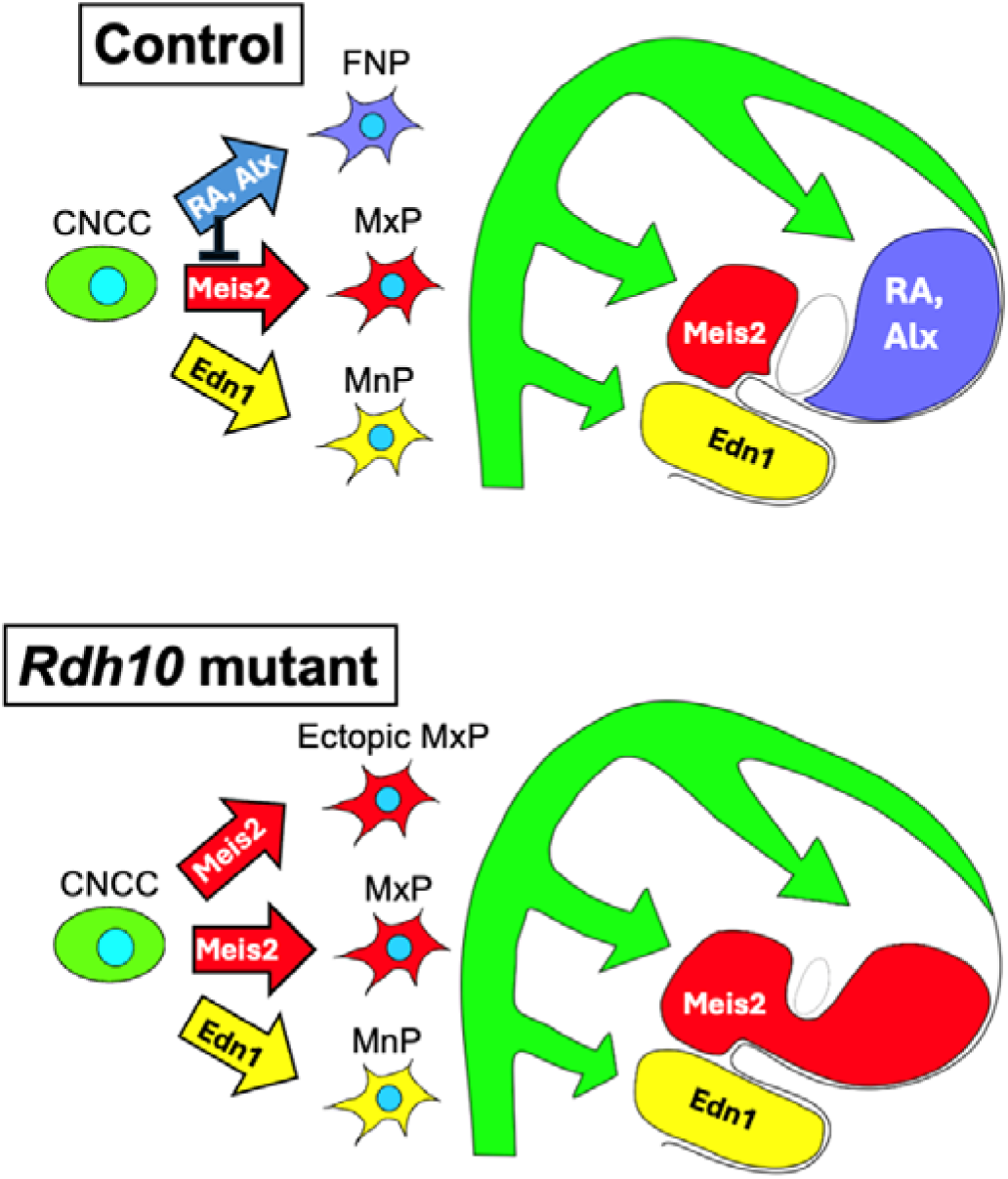
Schematic illustrating the molecular mechanisms governing FNP, MxP, and MnP identity. In control embryos, plastic CNCCs migrate ventrolaterally to distinct craniofacial prominences, where region-specific signaling cues establish positional identity. Within the FNP, retinoic acid signaling activates FNP-specific transcriptional programs, including Alx genes, while repressing maxillary determinants such as *Meis2*, thereby directing CNCCs toward a frontonasal fate. In the MxP, expression of maxillary transcription factors including *Meis2* promotes maxillary identity. In the MnP, *Edn1* induces mandibular-specific transcription factors such as *Dlx5/6* and *Hand2*, specifying mandibular fate. In *Rdh10* loss-of-function mutants, reduced retinoic acid synthesis in the FNP leads to derepression of maxillary transcription factors, including *Meis2*, resulting in ectopic activation of a maxillary gene regulatory network within the frontonasal mesenchyme. Consequently, CNCCs in the FNP undergo a fate transformation toward a maxillary identity. CNCC, cranial neural crest cell; FNP, frontonasal process; MxP, maxillary process; MnP, mandibular process.

## Materials and Methods

### Mice, and tamoxifen and all-trans retinoic acid administration

The mouse lines used in this study include *R26-CreER^T2^*(Jax stock #008463)(69), *Rdh10^fl/fl^* (21), *RARE-lacZ* (Jax stock #008477) (70, 71), Institute of Cancer Research (ICR) (CLEA, Japan). All mice were maintained under standard conditions as previously described (27). To generate conditional *Rdh10* knockout embryos, *R26-CreER^T2^;Rdh10^fl/fl^;RARE-lacZ* males were bred with *Rdh10^fl/fl^* females. Tamoxifen (5mg of tamoxifen and 1mg of progesterone in 100 μl of corn oil) was administered to pregnant females at E7.0 as previously described (27), to induce the deletion of *Rdh10* in Cre-positive embryos, while Cre-negative littermates were used as controls. To generate embryos with elevated retinoid signaling, pregnant ICR or *RARE-lacZ* females were administrated all-trans retinoic acid dissolved in the carrier by oral gavage, starting at E8.5 and continuing twice daily until the desired collection day. Control groups were administrated an equivalent volume of the carrier solution. For all-trans retinoic acid preparation, all-trans retinoic acid powder was dissolved in dimethylsulphoxide (DMSO) to a stock concentration of 25 mg/ml and then diluted 1:10 in corn oil just before use. The final administered volume was adjusted based on body weight to achieve a final dose of 25 mg/kg body weight.

### X-gal staining

To visualize retinoid signaling activity, *RARE-lacZ* embryos were dissected in cold PBS, then transferred to freshly prepared fixation solution (see below) on ice. Embryos older than E9.5 were fixed for 30 minutes, while those younger than E9.5 were fixed for 15 minutes. After fixation, embryos were washed three times with PBS and incubated in X-gal staining solution (see below) until the appropriate β-galactosidase intensity was observed. The staining reaction was stopped by washing the embryos in PBS. The fixation solution consisted of 2% formaldehyde and 0.2% glutaraldehyde, prepared in PBS. The staining solution was prepared in PBS and contained the following components: 1 mg/ml X-Gal, 5 mM potassium ferricyanide, 5 mM potassium ferrocyanide, 2 mM MgCI, 0.01% sodium deoxycholate, and 0.02% Nonidet P-40 (NP-40).

### Morphological analysis

Embryos were fixed in 4% paraformaldehyde at 4°C overnight after dissection. Whole mount nuclear fluorescent imaging technique was performed according to the established protocol (72, 73).

### Histological analysis

After capturing the morphological pictures, samples were dehydrated through a graded ethanol series (70%, 80% and 100%), embedded in paraffin and cut into 7 μm sections. Serial sections were used for hematoxylin and eosin staining following a standard protocol (74). Every section was examined to ensure the consistency and reproducibility of histological features.

### *In situ* hybridization

Whole-mount *in situ* hybridization was performed, followed by sectioning of stained embryos as previously described (1). All RNA riboprobes used in this study were synthesized based on sequences from the Allen Brain Atlas (https://mouse.brain-map.org/). At least three control and *Rdh10* mutant embryos were analyzed with each probe.

### Immunofluorescence

Fresh embryos were dissected in ice-cold PBS, fixed in 4% paraformaldehyde at 4°C for 2 hours, and dehydrated through a sucrose gradient (10%, 20%, and 30%). Subsequently, the samples were equilibrated by replacing half of the 30% sucrose solution with OCT compound, followed by full embedding in OCT compound. The embedded tissues were sectioned into 10 μm-thick slices using a cryostat, and the sections were collected onto slides. The frozen sections were air-dried on a 37°C plate for 10 minutes, followed by two washes in PBS to remove residual OCT compound. The sections were then blocked with 3% BSA in PBT (1% Tween-20 in PBS) for 1 hour at room temperature. After blocking, the samples were incubated with primary antibodies diluted in blocking solution (1:300) overnight at 4°C. Following incubation, the sections were washed three times with PBS and then incubated with Alexa Fluor-conjugated secondary antibodies (Invitrogen) diluted in blocking solution (1:300) for 1 hour at room temperature. After secondary antibody incubation, the sections were washed three times in PBS and counterstained with DAPI (1:500 in PBT) for 10 minutes at room temperature. The slides were washed twice with PBS, mounted with the fluorescence mounting medium (DAKO, S302380), and observed under a fluorescence microscope. The following antibodies used in this study included: mouse anti LHX6 (Santa Cruz, sc-271433), rabbit anti MEIS2 (Proteintech, 11550-1-AP), rabbit anti PRRX1 (Abcam, ab211292), rabbit anti PBX1 (Cell Signaling, 4342), rabbit anti SIX1 (Sigma, HPA001893), rabbit anti MECOM (Cell Signaling, 2593).

### Single-cell RNA sequencing

Tissues from E11.5 embryos were collected in three groups: FNP from three *Rdh10^fl/fl^* FNP from three *R26-CreER^T2^;Rdh10^fl/fl^* and FNP with MxP from three *Rdh10^fl/fl^*. After microdissection, tissues were dissociated into single cells as previously described (31). Then gene expression libraries were constructed and sequenced following the Chromium Next GEM Single Cell 3’ Reagent Kits User Guide, processed at the Research Institute for Microbial Diseases at Osaka University. Quality control, mapping, and count table generation were conducted via the CellRanger pipeline against the mouse mm10 genome. The output data from the CellRanger was analyzed using the Seurat v5.1.0 package (75) in RStudio v4.4.1 adhering to the standard workflow. Low quality cells (<300 genes and >10% mitochondrial genes) were filtered out, resulting in final analysis of 9424 (control FNP), 7932 (mutant FNP) and 11116 (control FNP with MxP) cells. Cell populations were annotated based on cluster-specific gene expression using published markers and in vivo staining. Statistical analysis of cell-type proportions between control and *Rdh10*_mutant samples was performed in R using a two-proportion Z-test, with p < 0.05 considered statistically significant. Differentially expressed genes between clusters were defined using the following criteria: log fold-change ≥ 0.5 and ≤ −0.5, adj. P < 0.01. Mesenchymal cell trajectories were reconstructed using Monocle3 v1.3.7 (76).

### Single-cell multi-omics

FNP tissues from five E11.5 *R26-CreER^T2^;Rdh10^fl/fl^* embryos and a comparable number of controls were flash-frozen in liquid nitrogen after dissection. Single nuclei were isolated following the manufacturer’s protocol (10X Genomics, Nuclei Isolation from Embryonic Mouse Brain Tissue for Single Cell Multiome ATAC + Gene Expression Sequencing, protocol CG00036_RevD). Multi-omic libraries for snRNAseq and snATACseq from the same barcoded single nuclei were generated and sequenced following the Chromium Next GEM Single Cell Multiome ATAC + Gene Expression User Guide at the Research Institute for Microbial Diseases at Osaka University. Sequencing reads were preprocessed and aligned to the mouse mm10 genome using the Cell Ranger ARC v2.0.0 pipeline. The raw outputs from CellRanger were analyzed using Seurat v5.1.0 (75) and Signac v1.13.0 (77) packages in RStudio v4.4.1, following the tutorial (WNN analysis of 10X Multiome, RNA and ATAC). Transcription factors significantly enriched in both RNA expression and chromatin accessibility at their motif regions were identified using the Wilcoxon rank-sum test from the presto package in R, with log fold-change > 0 and adjusted p-value < 0.05 for both modalities. The average AUC from RNA and motif accessibility rankings was used to prioritize key regulators.

## Acknowledgement

This study was supported in part by grants from the Ministry of Education, Science, Sports, and Culture of Japan (no. 19H03858 and 23K27803 to HK and no. 21K10159 for HO). We also appreciate for the support of Bioinformatics Research Unit, Graduate School of Dentistry, Osaka University for analyzing the dataset of single cell RNAseq. Research in the Trainor lab is supported by the Stowers Institute for Medical Research.

## Author Contributions

LX: Conception or design of the work; acquisition, analysis, and interpretation of data; drafting and critical revision of the manuscript for important intellectual content. YW, MO, TT, YD, QW, XN, DM, HO, TI, MY, and LLS: Acquisition, analysis, and interpretation of data; LLS also contributed to critical revision of the manuscript for important intellectual content. PAT and TY: Conception or design of the work; interpretation of data; critical revision of the manuscript for important intellectual content. HK: Conception or design of the work; interpretation of data; drafting and critical revision of the manuscript for important intellectual content.

---

**Figure S1:**
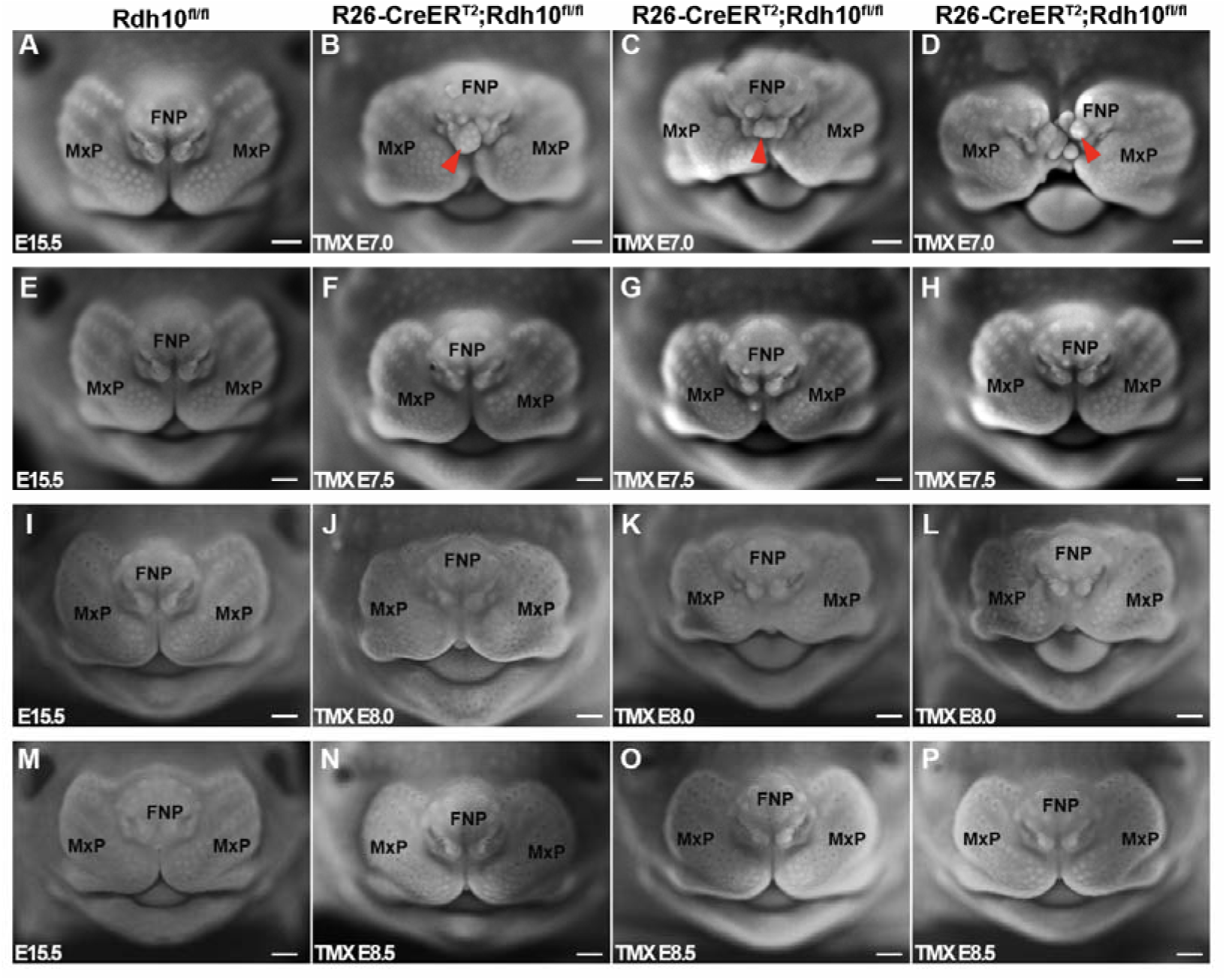
Temporal requirement of retinoid signaling during early CNCC specification. Tamoxifen was administered to pregnant females carrying *R26-CreER^T2^;Rdh10*^fl/fl^ embryos at different embryonic stages: E7.0 (A–D), E7.5 (E–H), E8.0 (I–L), or E8.5 (M–P). For each time point, the first panel shows a littermate control (A, E, I, and M, respectively), followed by three representative mutants (B–D, F–H, J–L, and N–P, respectively). Mutant embryos induced at E7.0 exhibit ectopic protrusions in the frontonasal region (red arrows). In contrast, embryos induced at E7.5, E8.0, or E8.5 do not display ectopic whisker pad formation at the stages examined. Scale bars, 500 μm. FNP, frontonasal process; MxP, maxillary process.

**Figure S2:**
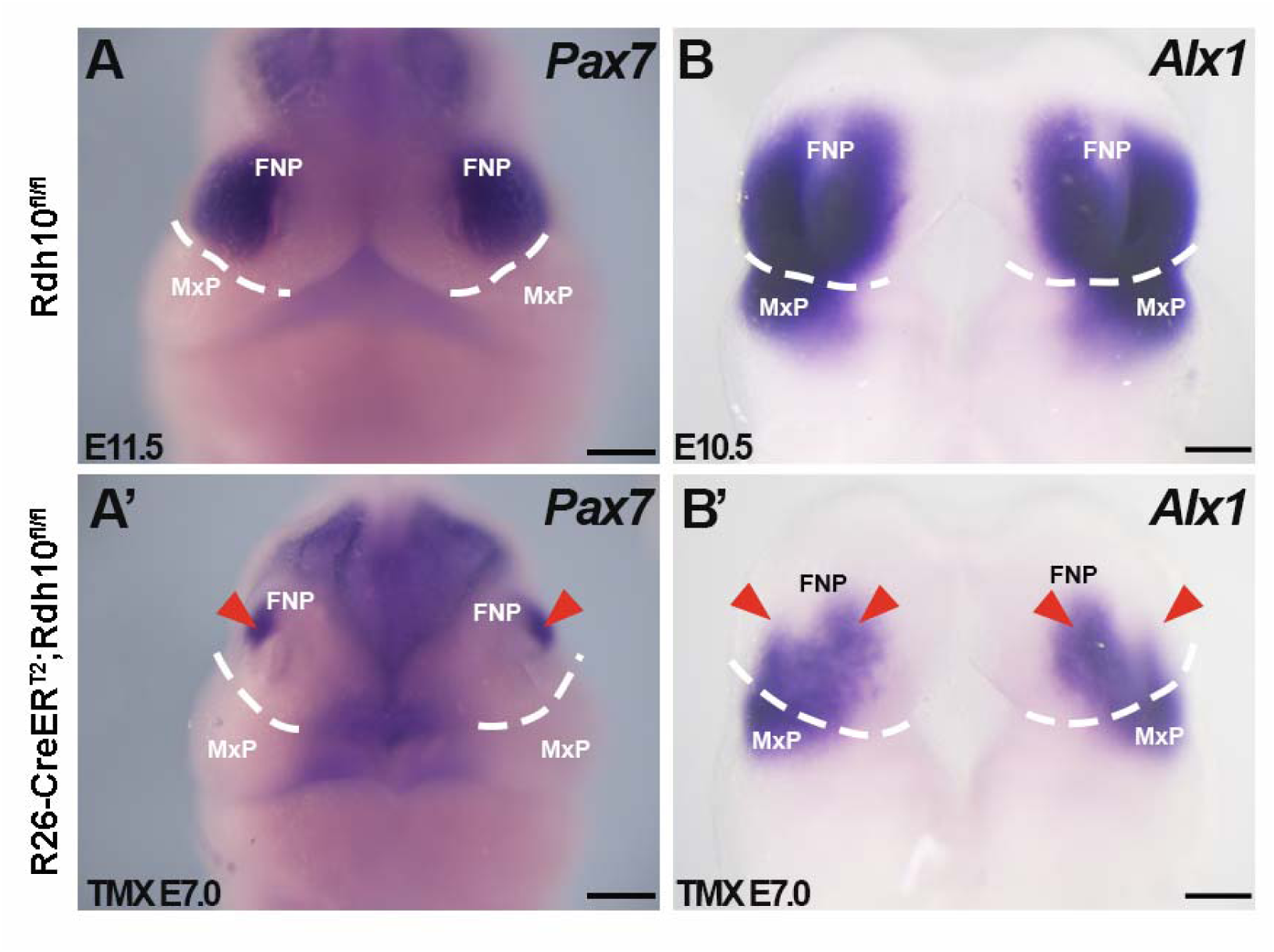
Diminished expression of frontonasal genes in *Rdh10* mutants. Frontal views of whole-mount in situ hybridization for *Pax7* and *Alx1* in control (A, B) and *Rdh10* mutant (A′, B′) embryos. The boundary between the FNP and MxP is outlined by a white dotted line. Red arrowheads indicate the reduced expression domain within the FNP region of mutants. Panels B and B′ were modified from previously published data (26). Scale bars, 500 μm (A, A’); 200 μm (B, B’). FNP, frontonasal process; MxP, maxillary process.

**Figure S3:**
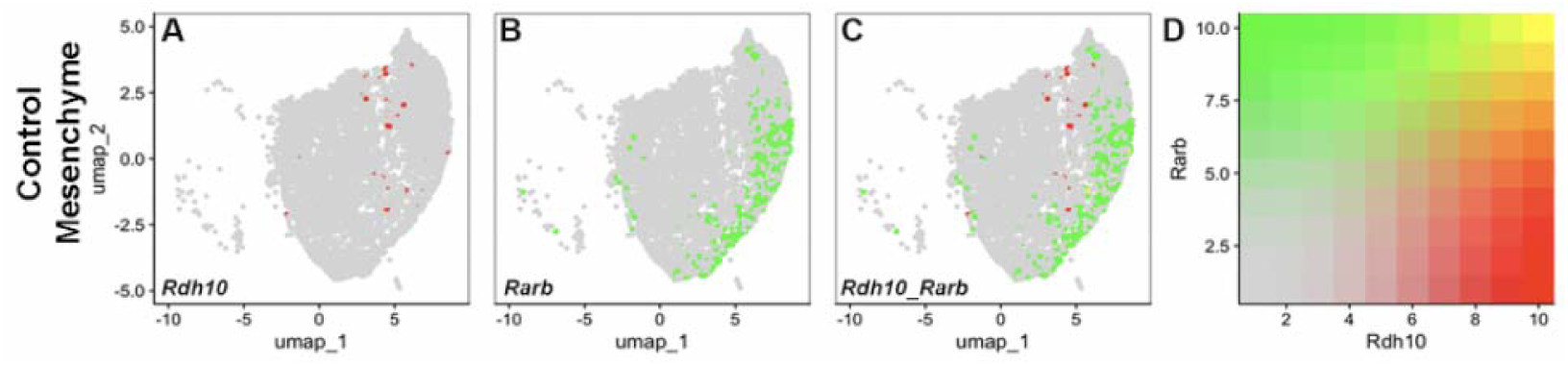
Visualization of retinoid signaling producing and receiving cells in control mesenchyme. (A–C) UMAP feature plots showing the expression of *Rdh10* (A) and *Rarb* (B), and blended visualization of *Rdh10* and *Rarb* co-expression (C) within the control mesenchymal population. In the blended plot, cells expressing *Rdh10* are shown in red, *Rarb* in green, and co-expressing cells in yellow. (D) Color scale indicating relative expression levels corresponding to the feature plots.

**Figure S4:**
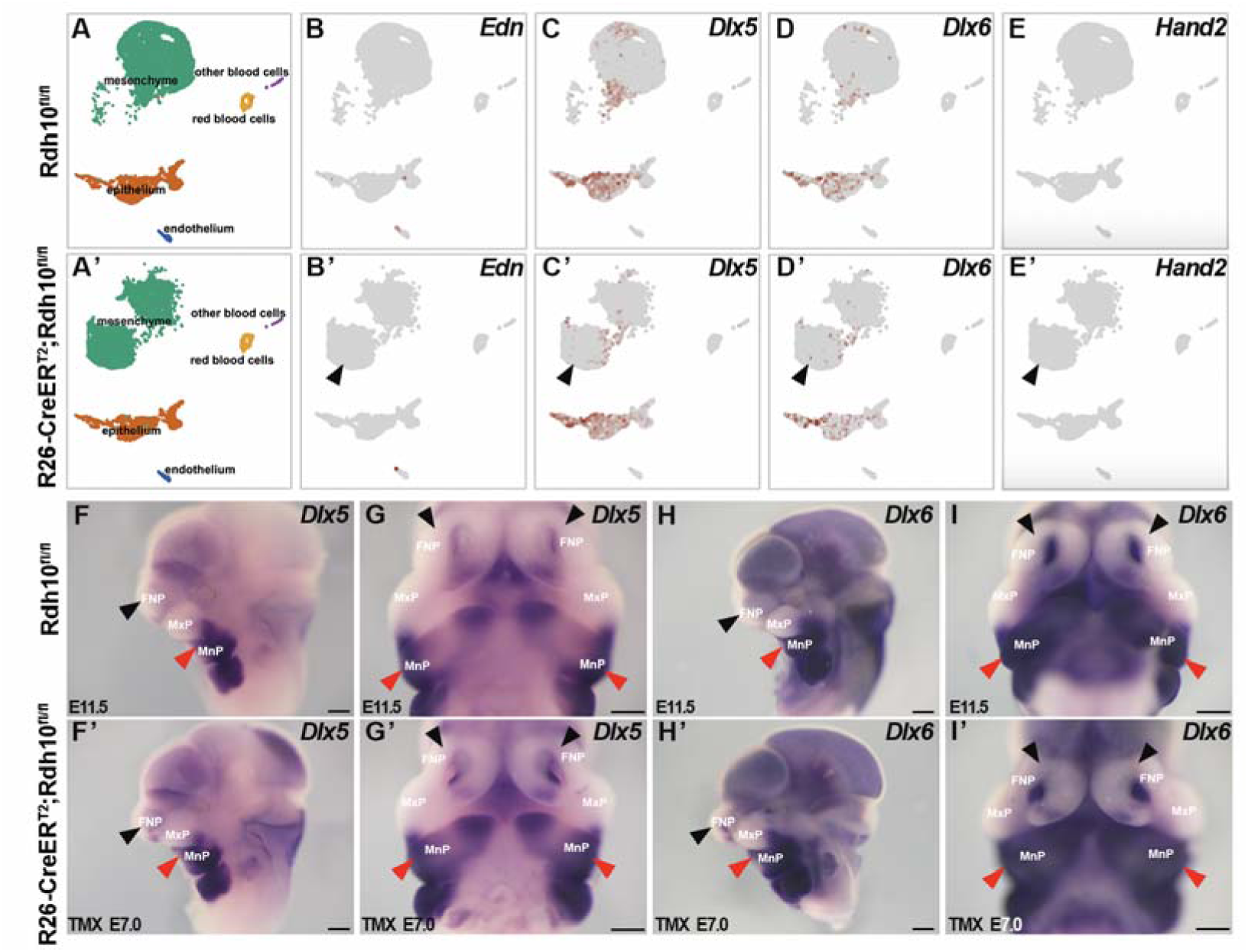
Lack of activation of the mandibular *Edn1–Dlx5/6–Hand2* signaling axis in retinoid signaling deficient mutants. (A, A′) UMAP visualization of control and *Rdh10* mutant samples. (B–E) Feature plots showing the expression of *Edn1, Dlx5, Dlx6,* and *Hand2* in control embryos. (B′–E′) Feature plots showing the expression of these genes in *Rdh10* mutant embryos. Black arrows in (B′–E′) indicate barely detectable expression in the ectopic mesenchymal cells of mutants. (F–G′) In vivo validation of *Dlx5* expression in control (F, G) and mutant (F′, G′) embryos. (H–I′) In vivo validation of *Dlx6* expression in control (H, I) and mutant (H′, I′) embryos. Panels (F, H, F′, H′) show lateral views, whereas (G, G′, I, I′) show frontal views. Black arrows in (F–I′) indicate minimal expression in the frontonasal region of both control and mutant embryos, whereas red arrows indicate strong expression in the mandibular region. Scale bars, 500 μm. FNP, frontonasal process; MxP, maxillary process; MnP, mandibular process.

**Figure S5:**
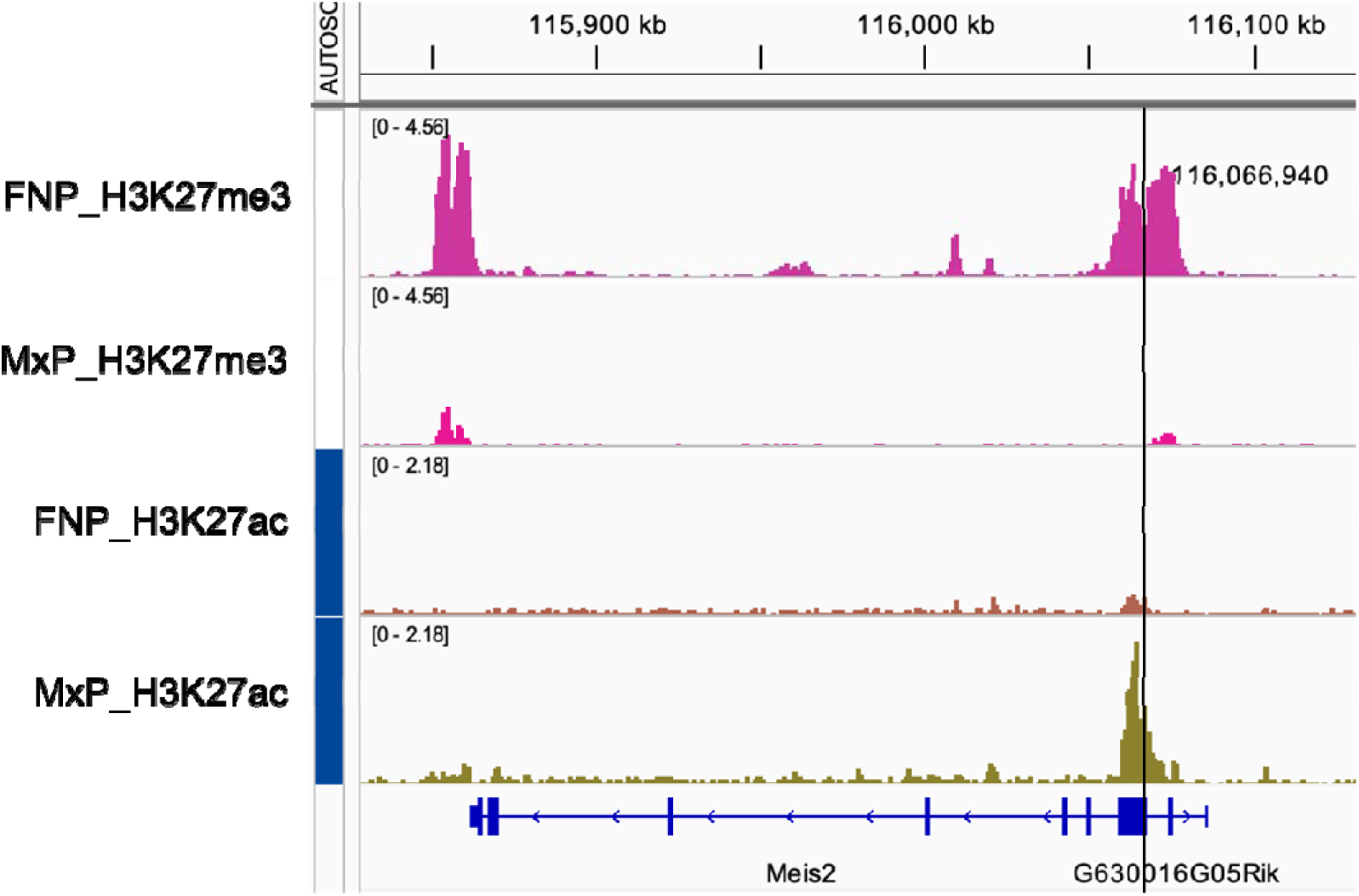
ChIP-seq analysis of histone modifications at *Meis2* locus. Re-analysis of published CHIP-seq data (36) to examine histone modifications associated with the active mark H3K27ac and the repressive mark H3K27me3 in cells isolated from the FNP and MxP of E10.5 wild-type embryos. The data reveal distinct histone modification patterns at the *Meis2* locus, with the dark line indicating its promoter region. FNP, frontonasal process; MxP, maxillary process.

**Table S1:**
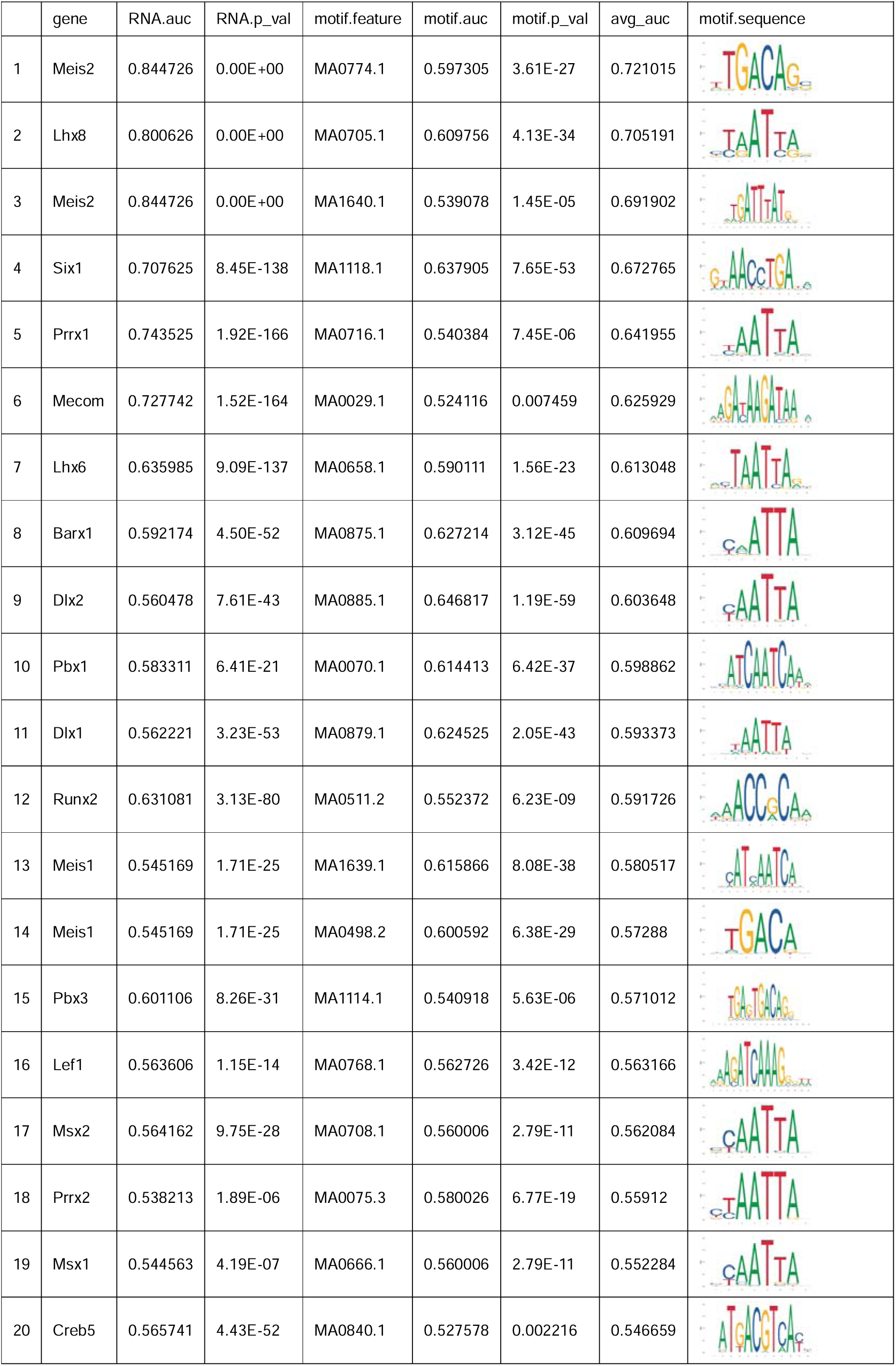
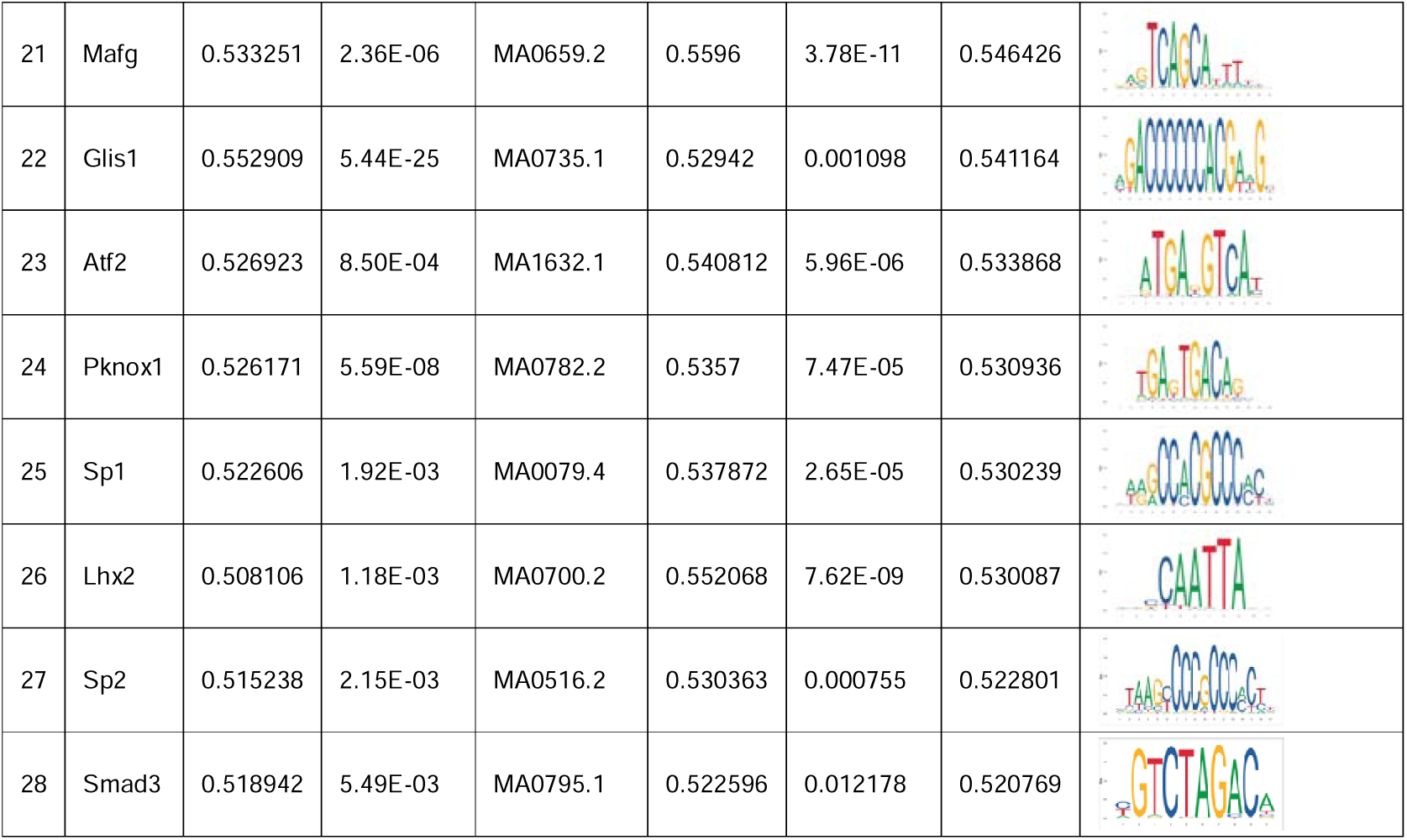
Single-cell multi-omics analysis identified 26 transcription factors.

