## Supplementary figures and images for "Retinoid signaling promotes frontonasal identity while repressing maxillary identity during craniofacial development"

### Supplemental Figure 1

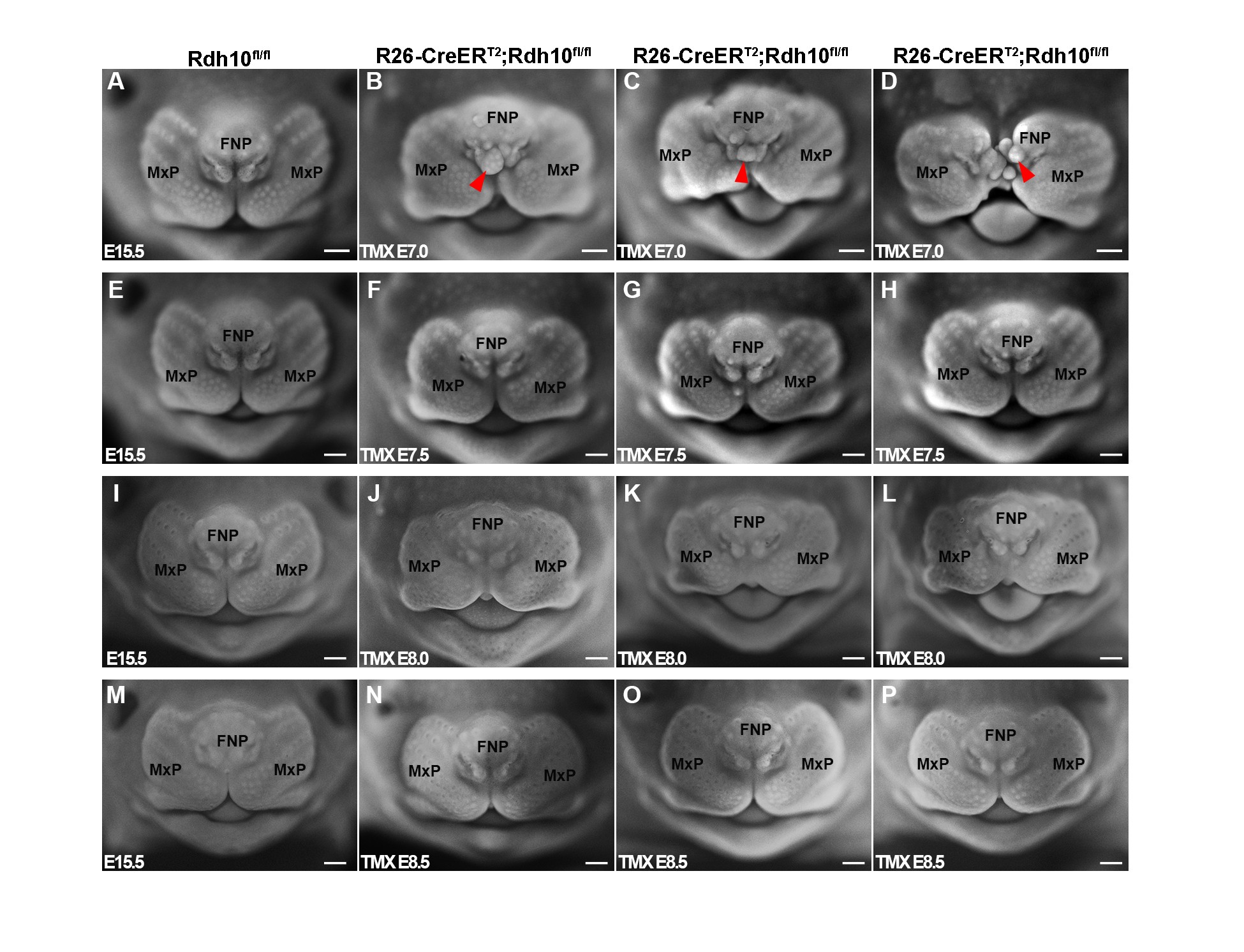

### Supplemental Figure 2

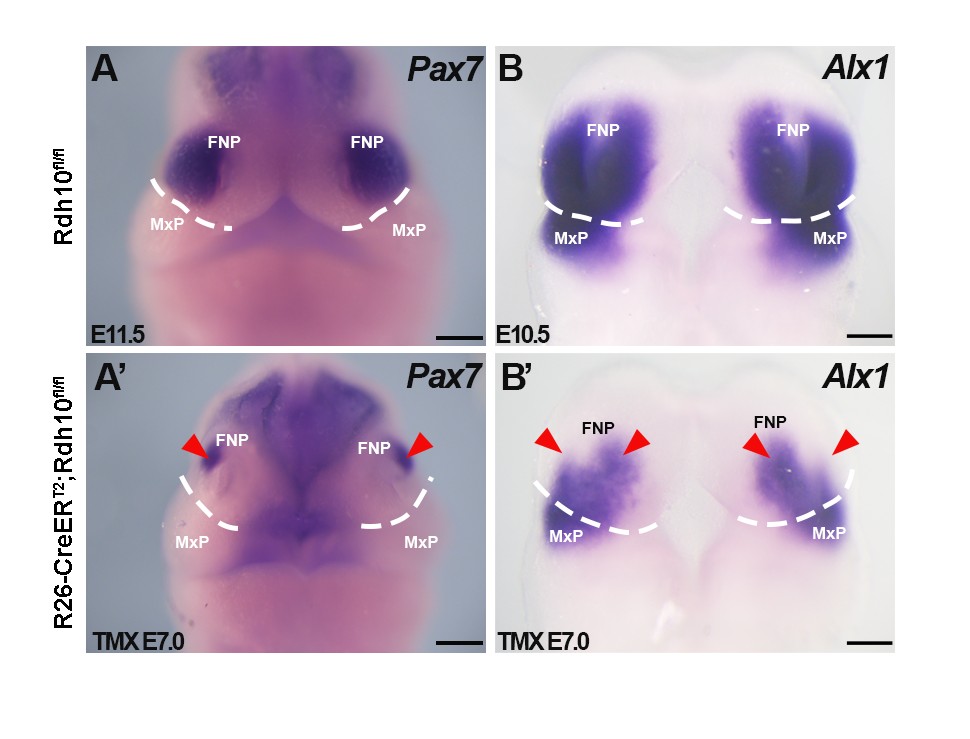

### Supplemental Figure 3

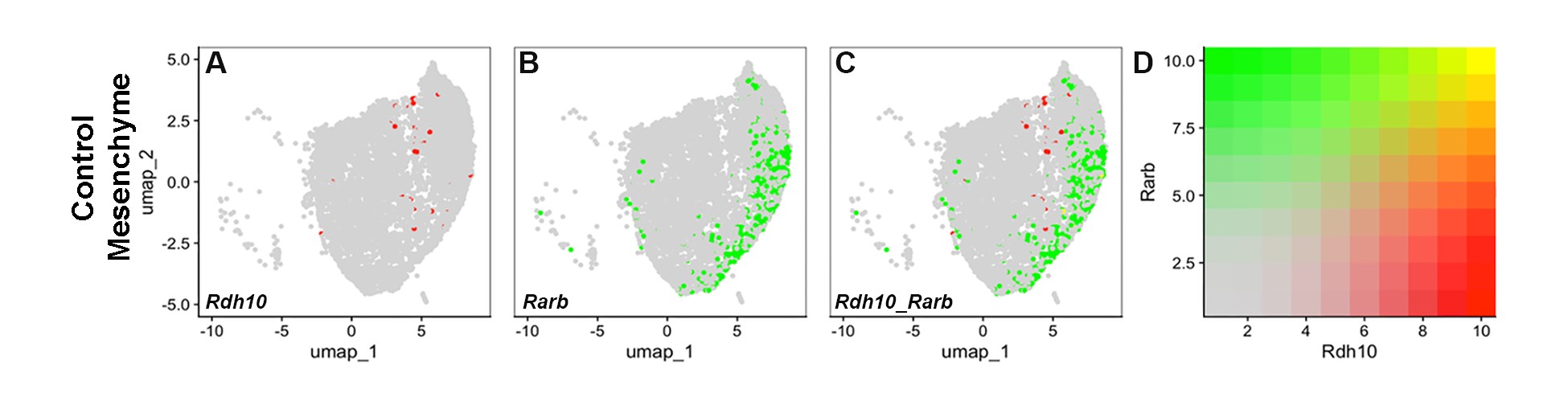

### Supplemental Figure 4

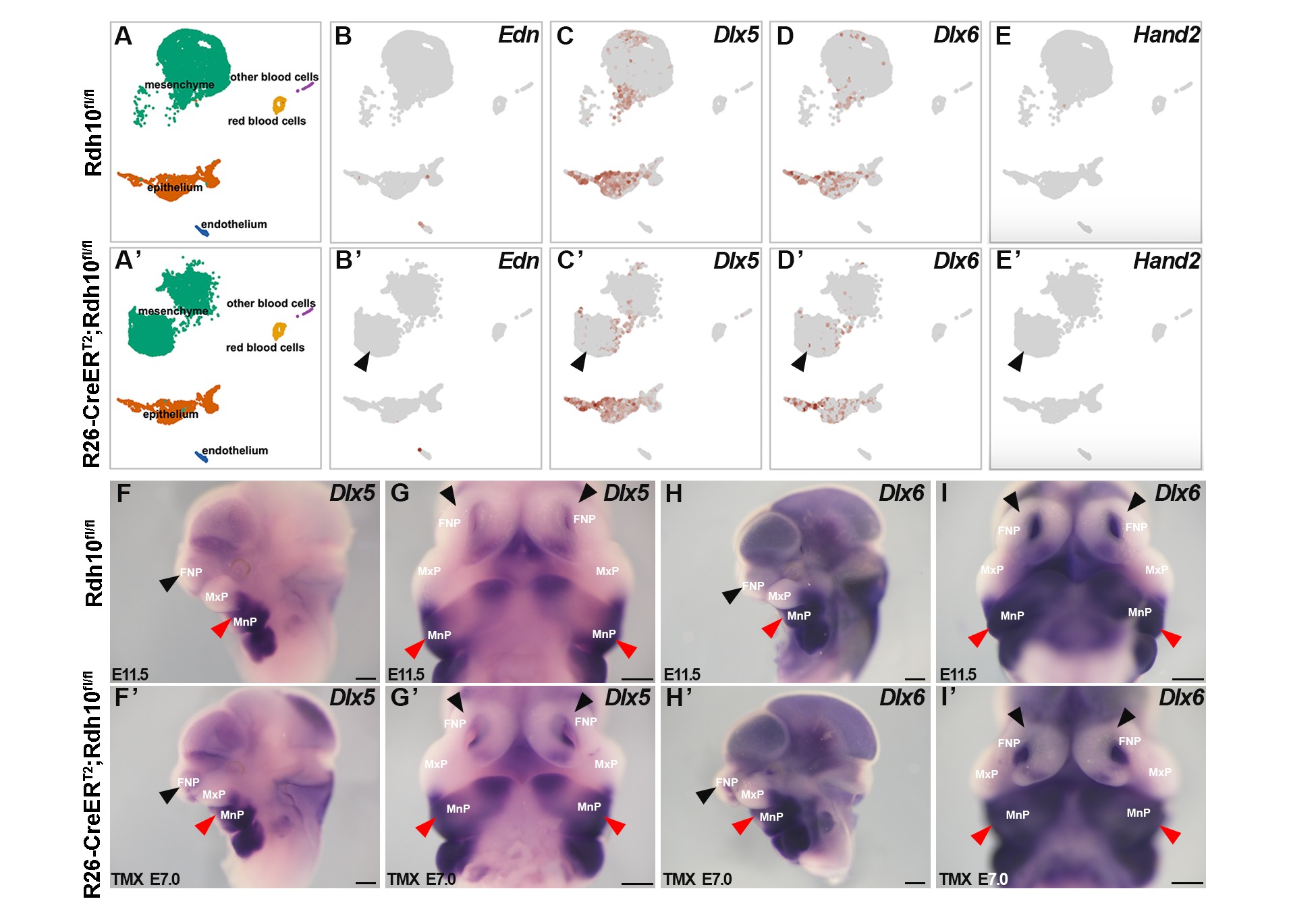

### Supplemental Figure 5

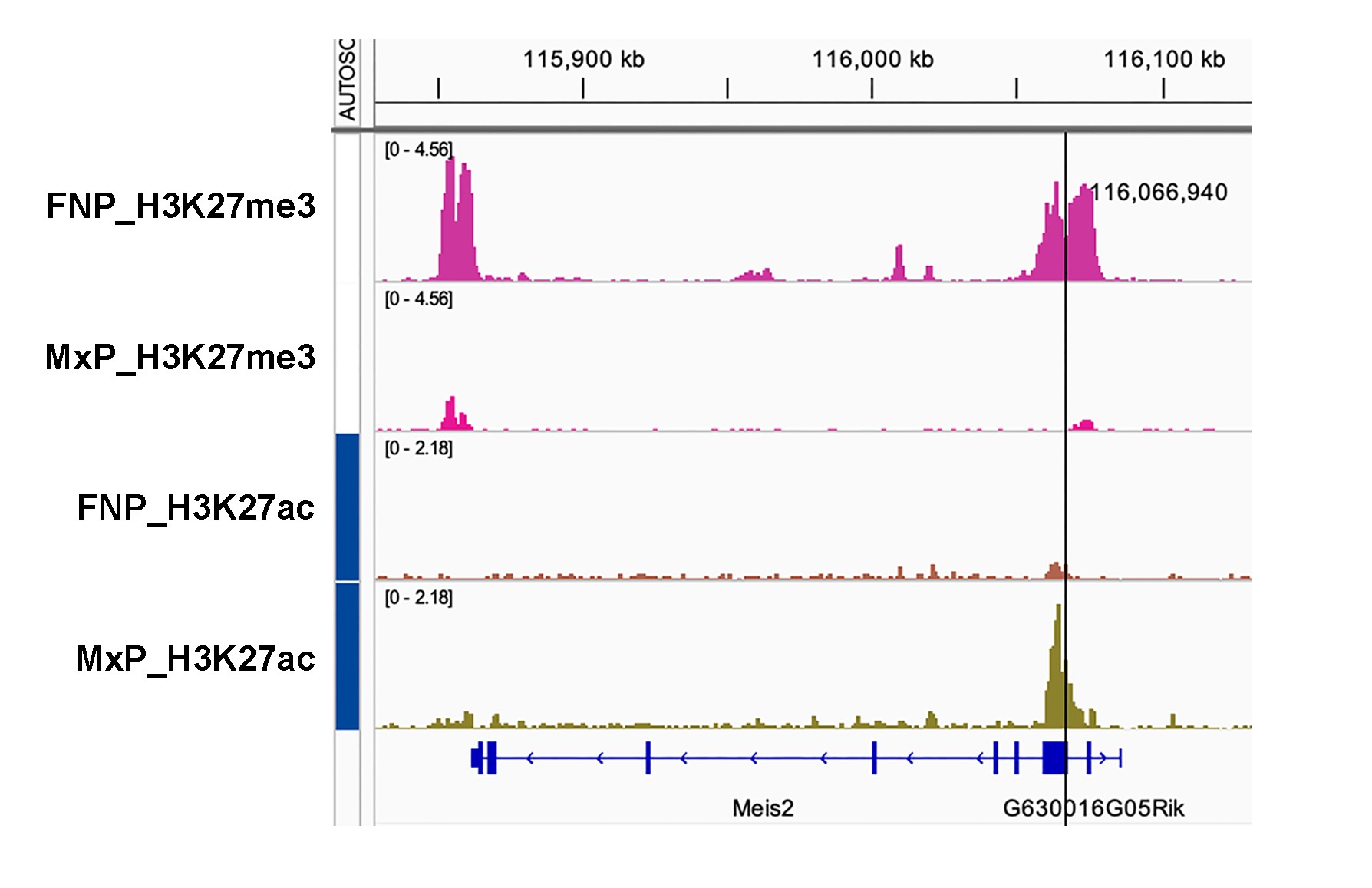
